# Peripheral heterochromatin tethering is required for chromatin-based nuclear mechanical response

**DOI:** 10.1101/2025.02.12.637704

**Authors:** A. Goktug Attar, Jaroslaw Paturej, Ozan S. Sariyer, Edward J. Banigan, Aykut Erbas

## Abstract

The cell nucleus is a mechanically responsive structure that governs how external forces affect chromosomes. Chromatin, particularly transcriptionally inactive heterochromatin, resists nuclear deformations through its mechanical response. However, chromatin also exhibits liquid-like properties, casting ambiguity on the physical mechanisms of chromatin-based nuclear elasticity. To determine how heterochromatin strengthens nuclear mechanical response, we performed polymer physics simulations of a nucleus model validated by micromechanical measurements and chromosome conformation capture data. The attachment of peripheral heterochromatin to the lamina is required to transmit forces directly to the chromatin and elicit its elastic response. Thus, increases in heterochromatin levels increase nuclear rigidity by increasing the linkages between chromatin and the lamina. Crosslinks within heterochromatin, such as HP1*α* proteins, can also stiffen nuclei, but only if chromatin is peripherally tethered. In contrast, heterochromatin affinity interactions that may drive liquid-liquid phase separation do not contribute to nuclear rigidity. When the nucleus is stretched, gel-like peripheral heterochromatin can bear stresses and deform, while the more fluid-like interior euchromatin is less perturbed. Thus, heterochromatin’s internal structure and stiffness may regulate nuclear mechanics via peripheral attachment to the lamina, while also enabling nuclear mechanosensing of external forces and external measurement of the nucleus’ internal architecture.

## I. INTRODUCTION

Cells are constantly subjected to mechanical stresses due to forces transmitted across whole tissue, individual cell interactions with the extra-cellular environment, constricted migration, internal cytoskeletal reorganization, and myriad other factors [1, 2]. Mechanical stresses deform nuclei [1–3], alter chromatin spatial organization [2, 4–7], modulate gene transcription [4, 8, 9], increase DNA damage [3, 10–14], transform or redistribute epigenetic marks [4, 5, 7, 15], and consequently, affect cell fate and identity [1, 3, 4]. These mechanobiological effects require the internal chromatin to sense the exterior of the nucleus or cell. In turn, they require physical mechanisms that couple disparate length scales such that mechanical forces acting at the nucleus scale (microns) can be propagated to the length scales of chromatin segments (tens of nanometers) [2, 16]. However, questions persist about the internal architecture and material state of chromatin [17], so the physical mechanisms underlying chromatin-based nuclear mechanical response remain unclear.

Experimental observations of chromatin’s material properties and physical organization suggest that chromatin can be liquid- and solid-like. *In vitro*, chromatin can exhibit properties of either state, depending on ionic and biochemical conditions [18, 19]. *In vivo*, single-nucleosomes [20, 21] and telomere-bound proteins [22] exhibit liquid-like dynamics, while viscoelastic measurements made by magnetically pulling a sub-micron chromatin locus [23] or inducing interfacial forces by condensation [24] suggest that chromatin may be described by a viscoelastic fluid. However, rheological measurements using half-micron magnetic beads suggest that chromatin is a polymer gel [25]. Furthermore, micromanipulation experiments stretching cell nuclei demonstrate that heterochromatin is a major contributor to the elastic response of the nucleus and similarly suggest gel-like chromatin [26–29]. These differing observations might be reconciled by the inhomogeneous chromatin elasticity observed in deformation microscopy [15, 30], optical tweezers [31], and localized thermal perturbation [32] experiments.

Heterochromatin has distinct features that enforce spatial inhomogeneity within the nucleus and contribute to nuclear mechanics. Heterochromatic regions are enriched in histone methylation and chromatin-crosslinking proteins, such as HP1*α*, both of which bolster nuclear stiffness [26–29, 33]. Furthermore, methylation and chromatin bridging by HP1*α* may drive spatial segregation by phase separation [34–36]. which may further strengthen chromatin and nuclei [37, 38]. Spatial segregation is further enforced in most cells by an array of proteins that peripherally tether chromatin to the nuclear lamina, including emerin, LBR, PRR14, and lamin A, among others [39–43]. In turn, heterochromatin localization at the periphery may protect the genome and facilitate nuclear mechanosensing by mechanically responding to applied stresses and stabilizing nuclear structure [44, 45].

Supporting this idea, cells dynamically alter heterochromatin levels, composition, and localization in response to various mechanical stimuli. In cyclically contracting cardiomyocytes, H3K9me3-marked heterochromatin accumulates at the nuclear periphery, while passive cardiomyocyte nuclei exhibit a uniform pattern of H3K9me3 [15]. Similarly, cell migration and associated nuclear deformations are associated with increases in H3K27me3 and peripheral H3K9me3 heterochromatin [5, 7, 14, 46–48]. In both cardiomyocytes and migrating cells, dense heterochromatic regions undergo larger deformations [15, 30] or changes in chromatin contacts [5, 6] than presumably softer euchromatic regions. Consistent with this protective effect, heterochromatin may form in sufficiently soft nuclei to inhibit DNA damage [3, 28]. However, in stiffer epidermal stem cell nuclei, applied forces have also been observed to reduce H3K9me3 heterochromatin, which decreases nuclear stiffness [3, 4]; this fluidizing effect inhibits DNA damage, perhaps because chromatin is less fragile in that scenario [2, 3]. Thus, heterochromatin is not only regulated by forces but also has a critical role in sensing, transmitting, and responding to them.

3D spatial modeling of chromatin suggests several possible mechanisms underlying different aspects of heterochromatin organization and structure. Polymer models with nonspecific affinity interactions between heterochromatic subunits and between heterochromatin and the lamina can capture compartmentalization and spatial localization of heterochromatin observed in chromosome conformation capture data (3C, Hi-C, Micro-C, etc.) and imaging experiments [49–52]. However, it was suggested that these models cannot explain nucleus stretching experiments [26]. Simulation models of chromatin as a uniformly crosslinked polymer gel linked by specific bonds to the nuclear lamina can recapitulate nuclear mechanical response [26, 29, 53], but are at odds with the nonuniform spatial distribution of heterochromatin [42] and the elasticity measured in chromatin locus pulling experiments [23]. Together, these models raise the questions of what internal structures enable heterochromatin levels to increase nuclear stiffness and how heterochromatin fulfills its apparent mechanical function.

To investigate the physical mechanisms by which heterochromatin contributes to nuclear stiffness and mechanosensing, we measure nuclear force response in a coarse-grained polymer physical model of the cell nucleus that includes epigenetic features of chromatin, chromatin-binding proteins, and a mechanically realistic nuclear lamina. We explored several scenarios: (i) heterochromatin that is unconstrained by specific linkages (*i*.*e*., neither internal crosslinking nor peripheral tethering to the lamina), (ii) heterochromatin that is constrained by internal crosslinks but not peripheral tethering, and (iii) heterochromatin that is tethered, with or without internal crosslinks. Our results revealed that heterochromatin enhances nuclear rigidity only when tethered to the nuclear lamina, facilitating the propagation of mechanical stress across the nucleus. Without chromatin tethering, heterochromatin failed to bolster nuclear stiffness even with strong crosslinking. Peripheral heterochromatin levels were directly linked to nuclear stiffness, with reduced heterochromatin tethering resulting in softer nuclei. Increasing heterochromatin levels further constrains heterochromatin through new lamina tethering and/or chromatin crosslinking, both of which increase nuclear stiffness. Additionally, we observed that lamina tethering and crosslinking direct mechanical deformations to the heterochromatin and lamina-associated domains (LADs). Our work thus uncovers a critical role for heterochromatin-lamina linkages in facilitating chromatin-based nuclear mechanical response and demonstrates how heterochromatin structure and mechanics be regulated to facilitate mechanosensing or protect the genome when nuclei are stressed and deformed.

## II. MATERIALS & METHODS

We performed molecular dynamics simulations of nuclei stretched by axially applied tensile forces using a polymer model of chromosomes inside of a polymeric lamina shell [26, 29, 53, 54]. Below, we briefly describe the simulation setup and procedure, and additional details are provided in the **Supplementary Information**.

### A. Chromatin polymer

Triblock copolymers model eight 240 Mb chromosomes. Each copolymer comprises *N* = 6002 monomeric subunits, where the two ends are telomeres, followed by 1000 subunits at one end that are designated as constitutive heterochromatin, and 5000 subunits representing genomic loci capable of forming A- and B-type compartments (euchromatin and facultative heterochromatin, respectively; **Fig. 1a**), which assigned based on chromosomes 1 and 2 of mouse rod cells (four of each) [52]. Subunits represent 40 kilobasepairs (kb) of chromatin, each of size *b* = 1*σ* ≈ 150 nm, where *σ* is the unit simulation length. Adjacent polymer subunits are connected by inextensible bond potentials. Interaction strengths between different chromatin types and between heterochromatin and shell subunits were swept and selected to model chromatin phase separation as in previous work [50, 52, 54, 55].

**FIG. 1.**
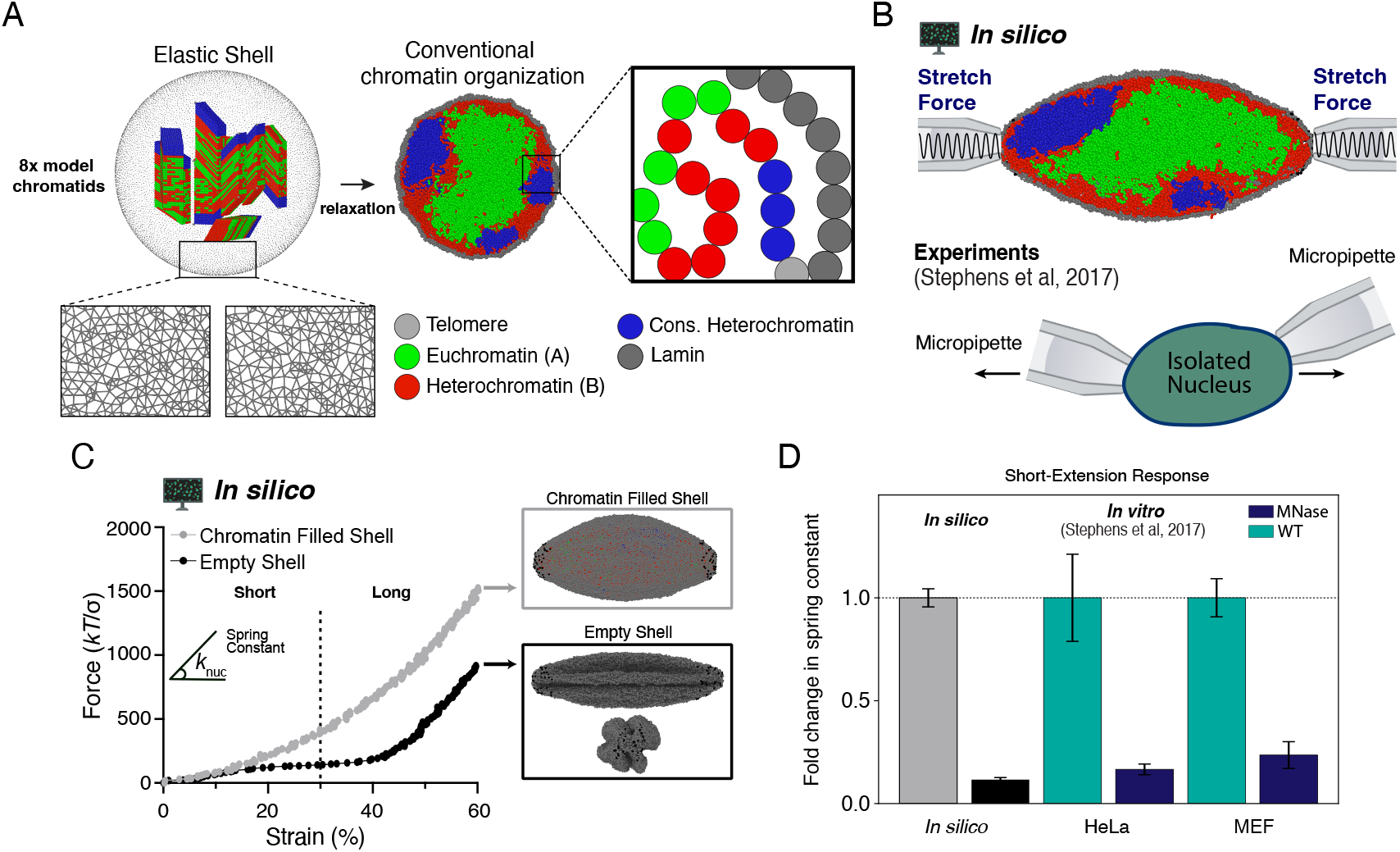
Heteropolymer simulation model recapitulates cell nuclear mechanical response. **A)** Eight chromatid-like structures are relaxed into a phase-separated mixture of constitutive and facultative heterochromatin (blue and red) and euchromatin (green) within a polymeric lamina (gray). **B)** Simulation snapshot with a schematic illustration of applied axial forces (top) and illustration of corresponding micromanipulation force spectroscopy experiment. **C)** Representative force-strain curves measured in simulations of a chromatin-filled nucleus (gray) and an empty polymeric shell (*i*.*e*., only the lamina; black). Insets show simulation snapshots of the deformed morphologies; empty shell inset also shows lamina buckles in a rotated view. Fold change in short-extension spring constant from wildtype or typical simulation nuclei (turquoise or gray, respectively) to MNase-treated nuclei or empty polymeric shells (navy blue or black) regimes in experiments [26] and simulations.

Chromatin polymers are initially compact, chromatid-like structures placed randomly near the nuclear center, and they are subsequently relaxed (**Fig. 1a**). Chromatin fills the spherical volume and segregates into heterochromatin-rich and poor domains, as expected [40, 42, 52]. In simulations with heterochromatin tethering and/or crosslinks, permanent (unbreakable) tethering to the shell and/or crosslinking are introduced after the last time step of equilibration. Crosslinks were assigned to subunits in contact after equilibration. Each heterochromatin subunit is crosslinked with probability *P*_c_ = 0.2 (unless noted) to a maximum of two others that are in contact (*i*.*e*., *r*_H_ ≤ 2.5*σ*). Chromatin subunits within *r*_L_ ≤ 2.5*σ* of the lamina were connected with a probability *P*_t_ to one lamina subunit, with *P*_t_ = 1.0 unless noted. For models of different levels or architectures of heterochromatin, perturbations to the control structure are introduced after the initial equilibration. In simulations altering heterochromatin fraction, *f*, only facultative heterochromatin is changed. Crosslinks and tethers are assigned after a change in the heterochromatin fractions, the numbers of tethers and crosslinks correlate with the alteration in facultative heterochromatin levels.

### B. Nuclear lamina shell

We modeled the lamina as an elastic, spherical polymeric shell using a bead-spring network of 15,000 monomeric subunits (**Fig. 1a**). This network mimics the nuclear lamina meshwork [56–58] and reproduces the stress-strain responses of isolated nuclei in micromanipulation experiments [26, 53, 59]. Lamina subunits were randomly positioned on the surface of a sphere with an initial radius *R*_*i*_ = 42*σ*. Subunits were connected by inextensible bond potentials, with each subunit forming 5 ≤ *n*_bond_ ≤ 8 bonds. After chromatin polymer relaxation, the shell radius is reduced to a starting radius of *R*_*s*_ = 32*σ*. After further relaxation of the composite polymer-shell structure, the shell relaxes to *R*_0_ ≈ 29*σ*, achieving a chromatin volume fraction of *ϕ* ≈ 23% [60] and a nuclear diameter of ≈ 10 *µ*m.

### C. Force spectroscopy simulations

Nuclei were stretched by applying axial pulling forces at two opposing poles as in micromanipulation experiments [26, 53] (**Fig. 1b**). To mimic partial aspiration by the micropipettes, 80 subunits from each pole (|*y*| ≥ 28*σ*) were randomly selected at the start of the simulation to be pulled (**Fig. S5**). Pulling was achieved by harmonic spring potentials, and forces exerted by these potentials were recorded. Springs with stiffness *K* = 20*k*_*B*_*T/σ*^2^ were initially positioned at *y*_0_ = ±60*σ* (with the nucleus centered at the origin), and each was moved away at constant velocity *v*_pull_ = 0.1*σ/τ*, where *τ* = 500 time steps (**Fig. 1b** and **S6**). The pulling force is given by *F* = ±*K*(*v*_pull_*t* − *y*_0_), where *t* is the elapsed time. No qualitative changes were observed with variations in pulling velocity or spring constant (**Fig. S6**). Each pulling simulation ran for 2 × 10^5^ time steps, resulting in strains up to *λ* ≡ Δ*L/R*_0_ ≈ 80%. The built-in *fix smd* algorithm within the LAMMPS MD package was utilized [61]. Ovito is used for visualization of trajectories [62]. Ten replicates were performed per simulation condition.

### D. Chromatin contact frequencies≤

To compute chromatin contact frequencies, as measured in Hi-C experiments [52, 63], we counted subunits within *r* ≤ *r*_*c*_ = 2.5*σ* at each simulation frame. *P* (*s*) curves were computed for facultative heterochromatin and/or euchromatin only. Similarly, chromatin contacts with the lamina were computed as a contact index, which was taken to be 1 for subunits within *r*_*L*_ ≤ 2.5*σ* of the lamina and 0 otherwise. Measurements were averaged over 10 simulation replicates, using 10 configurations (snapshots) from each simulation just before stretching to 30% strain, and unless noted, 110 configurations just after stretching.

## III. RESULTS

### A. Chromatin governs nuclear force response to small to moderate deformations

Previous force spectroscopy experiments with isolated nuclei established the distinct mechanical contribution of chromatin to nuclear rigidity, independent of lamin proteins [26, 27, 29, 31, 64]. We first used force spectroscopy simulations to assess whether our multiblock copolymer simulation model captures the contribution of chromatin to nuclear stiffness.

Simulated nuclei were progressively stretched via forces applied to the lamin subunits at opposite poles (**Fig. 1c**; see Materials and Methods). Forces were applied to lamins by spring potentials whose centers were moved apart at a constant velocity, mimicking the experimental pulling protocol [26]. Stretching simulations of a chromatin-filled polymeric shell produced a strain-stiffening force response, with the measured force increasing more abruptly for larger strains (**Fig. 1c**, gray), as in experiments [26, 31]. Thus, our model is consistent with nuclear force response to dynamic stretching.

To confirm that chromatin is essential for nuclear stiffness at small deformations, we stretched empty polymeric shells (*i*.*e*., without the chromatin polymer). These simulations modeled treatment by the enzyme MNase, which digests chromatin. The empty shell produced a minuscule force response, with an effective spring constant (local slope of the force-extension curve), *k*_nuc_, almost an order of magnitude lower than that of the chromatin-filled nucleus. This behavior persisted up to strains of ~ 40% (**Fig. 1c**, black), in agreement with previous experiments and simulations [26, 31, 53]. Beyond 40% strain, deformation to the lamina increases the stress and alignment in lamin-lamin bonds along the direction of the applied force [26, 53] (**Fig. S4**). Deformed empty nuclei have smaller cross-sectional dimensions, shrink, and exhibit longitudinal buckles (**Fig. 1c**), as in previous simulations and experiments [53]. Therefore, the chromatin polymer is critical for robust mechanical response to small strains.

### B. Peripheral localization and phase separation of heterochromatin alone do not increase nuclear rigidity

Heterochromatin is paramount to chromatin-based nuclear rigidity [3, 26, 27, 29, 33]. Heterochromatin is thought to be established, maintained, and peripherally localized by polymer phase segregation [19, 34, 35, 49, 50, 52, 55, 65–69] and interactions with the nuclear lamina [39, 40, 43, 52, 70, 71]. Therefore, we investigated how alterations to heterochromatin phase separation impact nuclear stiffness.

We tested whether increasing the prevalence of the heterochromatic phase would increase nuclear stiffness, similar to experimental observations [26, 27, 29]. We randomly converted heterochromatin subunits into euchromatin or vice versa to tune total heterochromatin content to a prescribed level (**Fig. S8**). This procedure also altered the peripheral localization of heterochromatin because heterochromatin subunits have a nonspecific affinity to the lamina [52, 54]. Surprisingly, we found that nuclear force response does not increase with increasing heterochromatin content (**Fig. 2a-c**), but rather, slightly *decreases*. This is due to the larger polymer osmotic pressure of the less compact euchromatin phase [72]. This reinforces the notion that the origin of chromatin-based stiffness in cell nuclei is not osmotic pressure alone [26]. Together, our findings suggest that heterochromatin phase separation and peripheral localization do not govern nuclear rigidity on their own.

**FIG. 2.**
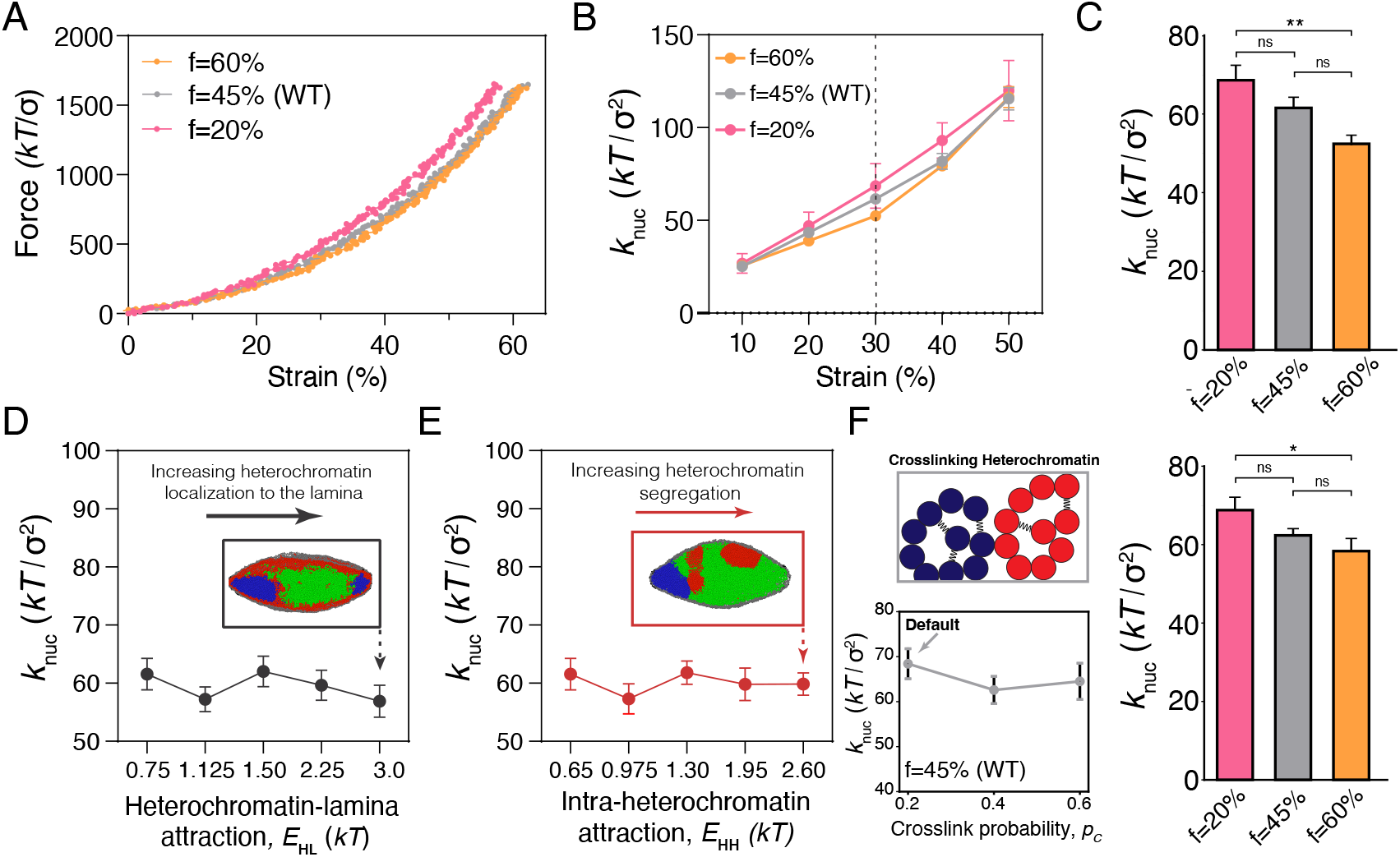
Effects of heterochromatin microphase separation and internal structure on nuclear mechanical response. **A)** Representative force-strain curves for nuclei with different fractions, *f*, of facultative heterochromatin, in nuclei with nonspecific affinity interactions between heterochromatin and the lamina. **B)** The nuclear spring constant averaged over ten replicates for various facultative heterochromatin levels, *f*, at different strains. **C)** Nuclear spring constants computed from the slope of the force-strain curves at 30% strain. ** indicates *p <* 0.01, while ns denotes not significant. **D)** Nuclear spring constants at 30% strain as a function of the heterochromatin-lamina subunit attraction energy, *E*HL and **E)** the intra-heterochromatin subunit attraction energy, *E*_HH_. Inset images show representative snapshots of nuclear deformations and spatial organization of euchromatin (green), facultative heterochromatin (red), and constitutive heterochromatin (blue) contained within the nuclear lamina (gray). **F)** *Top left* : schematic illustration of crosslinking within heterochromatin. Constitutive (blue) and facultative (red) heterochromatin may crosslink to other heterochromatin subunits of the same type. *Bottom left* : spring constants, *k*_nuc_ at 30% strain for different levels of crosslinking, set by crosslink probability *p*_*c*_. *Right* : bar plot showing spring constants versus heterochromatin fraction, *f*, with *p*_*c*_ = 0.2 such that the total number of crosslinks increases with *f*. Data are shown as mean ±SEM. Statistical significance is assessed by one-way ANOVA followed by a post-hoc Tukey’s HSD test for pairwise comparisons. * denotes *p <* 0.05, ** denotes *p <* 0.01, and *** denotes *p <* 0.001.

Since heterochromatin should strengthen nuclear force response, we speculated that the heterochromatin affinity interactions in our simulations might be too weak. However, despite increasing the heterochromatin-lamina affinity fourfold, the effective nuclear spring constant, *k*_nuc_, did not significantly change (**Fig. 2d**). Furthermore, *k*_nuc_ remained insensitive to alterations in heterochromatin levels (**Fig. S7**). Similarly, heterochromatin self-affinity did not affect the nuclear spring constants for a broad range of affinities (**Fig. 2e**). Even in simulations with strong affinity interactions, the heterochromatin phase could move along the nuclear periphery (*e*.*g*., inset to **Fig. 2d**) and/or internally rearrange as the nucleus deformed, without altering the number, and thus, the total energy, of heterochromatin-lamina and heterochromatin-heterochromatin contacts. We conclude that while heterochromatin-lamina attractions and heterochromatin self-affinity may govern mesoscale chromatin phase separation, they do not provide a robust mechanical coupling between nuclear deformation and heterochromatin and, therefore, do not promote nuclear rigidity.

### C. Heterochromatin-based nuclear stiffness cannot be explained by chromatin crosslinking alone

We next turned to a mechanism previously shown to bolster chromatin-based nuclear rigidity in both experiments and simulations: chromatin-chromatin crosslinking [29, 37]. To model crosslinks, such as HP1*α* [73, 74], *we introduced permanent bonds between heterochromatin subunits. Surprisingly, heterochromatin crosslinking slightly decreased* nuclear stiffness in force spectroscopy simulations, similar to the effects of affinity interactions (**Fig. 2f**). This shows that externally applied forces that generate small deformations (*<* 30% strain) do not deform the chromatin segments constrained by crosslinks, so crosslinks do not increase nuclear rigidity in this scenario. Therefore, crosslinks alone do not explain the chromatin-based nuclear mechanical response. This result suggests that the model lacks a key mechanical component required to generate force response.

### D. Tethering heterochromatin to the lamina is required for heterochromatin-driven nuclear force response

We sought to understand why alterations to heterochromatin polymer structure and organization did not elicit a strong change in force response (**Fig. 2**). In our initial model, interactions between heterochromatin and the lamina were nonspecific, allowing significant chromatin mobility. However, in contrast, many cells have specific tethers between chromatin and the lamina or nuclear envelope, either through lamin A itself or lamina- or envelope-associated proteins, including LBR, emerin, Lap2, and PRR14, among others [40, 41, 43, 70, 75, 76]. Therefore, we tested whether specifically tethering heterochromatin to the lamina would mechanically couple heterochromatin architecture to nuclear force response.

We incorporated specific, unbreakable, stretchable heterochromatin-lamina tethers into the model (**Fig. 3a**; see Materials and Methods) and simulated nuclear stretching. Nuclei with chromatin-lamina tethers exhibited a systematic increase in nuclear stiffness for small to moderate deformations (**Fig. 3b-d**), consistent with previous research [29, 75]; the effective nuclear spring constant for small deformations is twofold higher in nuclei with chromatin tethering than those without. Furthermore, in simulations with moderate amounts of heterochromatin tethering (*P*_*t*_ *<* 1), nuclei exhibited an intermediate force response (**Fig. 3d**). Visual inspection reveals that the bulk spatial organization of the chromatin phases after deformation is similar in both cases (**Figs. 3c**, inset). These observations demonstrate that the mechanical coupling imposed by heterochromatin tethers, but not peripheral heterochromatin localization alone, stiffens nuclei.

**FIG. 3.**
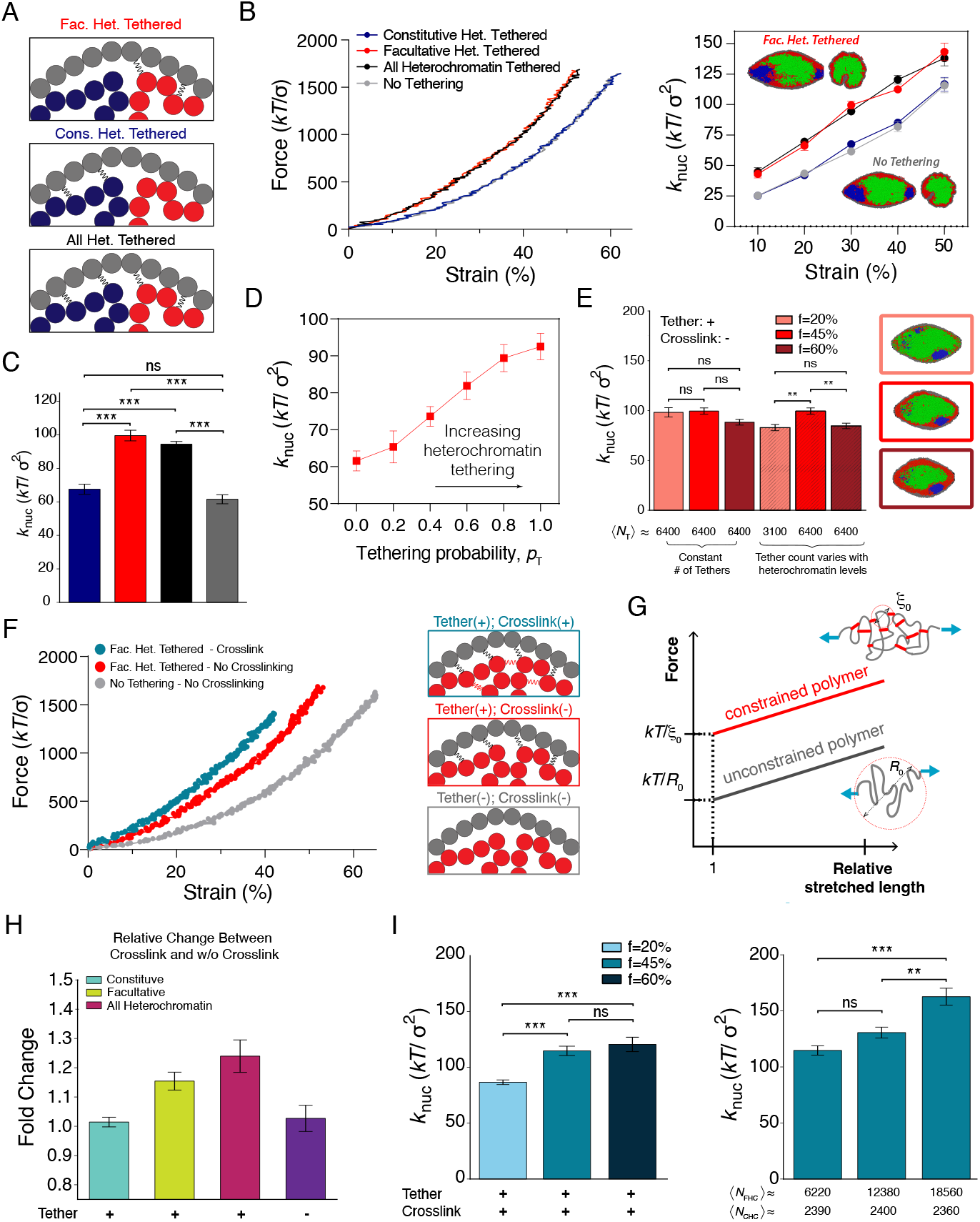
Nuclear mechanical effects of heterochromatin tethering and changes in tethered heterochromatin organization. **A)** Illustrations of specific tethering of heterochromatin to the lamina. **B)** Effective spring constants, *k*_nuc_, at 30% strain as a function of strain, obtained from at least 10 independent force spectroscopy simulations. **C)** *Left* : Representative force-strain curves for nuclei with and without tethering of peripheral heterochromatin to the lamina. *Right* : Spring constants, *k*_nuc_, as a function of strain. **D)** Nuclear spring constant as a function of facultative heterochromatin tethering probability, *p*_*T*_. **E)** Nuclear spring constant different levels of heterochromatin, *f*, where *f* =45% is the control case. Mean number of tethers, ⟨*N*_*T*_⟩, is either increased probabilistically (see Methods) with *f* or held constant, as indicated by the numbers below each bar. Simulations shown for tethering probability. **F)** *Left* : Representative force-strain curves for nuclei with and without heterochromatin crosslinking in the presence or absence of facultative heterochromatin tethering. *Right* : Illustrations of these scenarios. **G)** Schematic force-strain plot and illustration of expected scaling of force response as a function of relative stretched polymer length. Heterochromatin crosslinks act as constraints that decrease the length of stretchable polymer segments. **H)** Fold change in effective nuclear spring constant from untethered and uncrosslinked scenario (purple) for nuclei with specific tethering and crosslinking only within different types of heterochromatin: constitutive (turquoise), facultative (yellow), all (red). **I)** Effective spring constants with different levels of crosslinked heterochromatin with lamina tethering. *Left* : *k*_nuc_ with tethered and crosslinked heterochromatin for different heterochromatin fractions, *f*, with crosslinking probability *p*_*c*_ = 0.2 such that the number of crosslinks increases with *f*. *Right* : *k*_nuc_ with tethered and crosslinked heterochromatin at fixed *f*, but with crosslinking levels determined by different crosslinking probabilities, *p*_*c*_. Mean quantities of facultative and constitutive heterochromatin crosslinks, ⟨*N*_FHC_⟩ and ⟨*N*_CHC_⟩, respectively, are listed beneath each bar. Bars are mean ± SEM. Statistical significance is assessed by one-way ANOVA followed by a post-hoc Tukey’s HSD test for pairwise comparisons. * denotes *p <* 0.05, ** denotes *p <* 0.01, and *** denotes *p <* 0.001.

Between the two types of tethers linking heterochromatin to the lamina, those associated with facultative heterochromatin are the main contributors to nuclear rigidity (**Fig. 3a-c**). The facultative heterochromatin phase has a larger interaction area with the lamina, so, it has a larger number of tethers and mechanically couples the entire lamina. In contrast, tethering only constitutive heterochromatin (**Fig. 3a**) to the shell does not increase the nuclear spring constant (**Fig. 3b-c**). Therefore, although constitutive heterochromatin more strongly phase separates and peripherally localizes in the model [52, 54], it contributes less to nuclear mechanics. Together, our results indicate that one mechanism for heterochromatin-induced nuclear stiffness is the increase in chromatin-lamina tethering, which might coincide with expanding the lamina-associated heterochromatin phase.

### E. Chromatin-lamina tethering links heterochromatin levels to nuclear rigidity

To test whether linkages between chromatin and the lamina also directly connect nuclear mechanical properties and heterochromatin levels, we repeated the force spectroscopy simulation measurements on nuclei with specific heterochromatin-lamina linkages. With tethering to the lamina, reducing heterochromatin levels resulted in a systematic decrease in nuclear rigidity compared to the control case with “normal” heterochromatin levels (**Fig. 3e**). Increasing heterochromatin also resulted in a slight reduction in *k*_nuc_. Nonetheless, in contrast to the model without chromatin tethering (**Fig. 2a-c**), a ~ 25% decrease in heterochromatin can explain the reported decrease in the nuclear spring constant induced by inhibition of heterochromatic epigenetic modifications by methyltransferases in experiments [26–29] (**Fig. 3e**).

Heterochromatin-driven changes in nuclear stiffness were due to two factors: 1) chromatin condensation altering polymer osmotic pressure (as in **Fig. 2c**) and 2) changes in the number of tethers to the nuclear lamina. Decreasing heterochromatin fraction reduces the number of chromatin-lamina tethers, which decreases nuclear stiffness, despite a concurrent increase in polymer osmotic pressure due to chromatin decondensation. However, increasing heterochromatin fraction from *f* = 45% to *f* = 60% also slightly decreases *k*_nuc_ because the peripheral tethering is already saturated at *f* = 45%, but chromatin further condenses with this change. Consistent with this idea, if the total number of tethers is held constant, the model reproduces the trend observed without tethering (**Fig. 2c**): heterochromatin increases reduce nuclear stiffness (**Fig. 3e**).

Once again, these results indicate that nonspecific interactions between nuclear components are insufficient to produce the experimentally observed changes to nuclear rigidity. Instead, as shown by the stretching of chromosomal segments between tethered sites, it is the physical constraints of specific sites to the lamina that contribute to nuclear rigidity (**Fig. S13**). Therefore, the physical bridging of heterochromatin to the lamina provides the mechanical coupling needed to enable heterochromatin to respond to forces applied to the nucleus. Heterochromatin-driven nuclear stiffening can arise indirectly through heterochromatin’s effect on the quantity of chromatin-to-lamina linkages.

### F. Heterochromatin crosslinks directly contribute to nuclear force response when chromatin is tethered to the lamina

To determine whether heterochromatin can regulate nuclear stiffness through its own biophysical properties, we considered a model in which heterochromatin has internal crosslinks in addition to specific tethers to the nuclear lamina (**Fig. 3f, g**). In the presence of chromatin-lamina tethers, nuclear rigidity is sensitive to the degree of heterochromatin crosslinking, unlike simulations without tethers (**Fig. 2f-g,i**). With tethering, nuclei stiffen with increasing levels of heterochromatin crosslinking, even when the number of chromatin-lamina tethers is held constant (**Fig. 3h-i**). This is consistent with previous simulations with crosslinks throughout all chromatin [29]. Furthermore, with tethering and crosslinking, elevating the amount of heterochromatin can stiffen the nucleus because this perturbation also increases the total level of intra-chromatin crosslinking (**Fig. 3i**).

These observations sharply contrast with results for simulations without chromatin-lamina tethering (**Fig. 2f**). Therefore, while heterochromatin crosslinking may provide an important component of nuclear mechanical response, the effect depends on tethering to the lamina. Altogether, chromatin tethering to the lamina enables nuclear mechanosensing by providing the critical links for chromatin to “sense’ applied forces and external factors to “see” the internal physical organization and structure of the nucleus.

### G. Nuclear stretching increases lamina-associated heterochromatin

Our simulations indicate that mechanical communication between the nuclear periphery and interior arises due to physical constraints imposed by chromatin-lamina tethers. We next asked how these constraints affect force transmission into the nuclear interior and reorganize chromatin. To address this question, we performed *in silico* versions of DamID [77, 78] and Hi-C [63] by computing lamina-chromatin contacts and chromatin contact maps before and after nuclear stretching.

These analyses reveal small, but systematic, differences in peripheral heterochromatin rearrangement in nuclei with and without tethering to the lamina. With heterochromatin tethered to the lamina, nuclear deformations result in an expansion of lamina-associated domains (LADs), as measured by heterochromatin-lamina contacts (**Fig. 4a-b**). These new LADs form at the expense of chromatin-chromatin contacts, as shown by a systematic change in contact frequency (**Fig. S13**). In contrast, without peripheral heterochromatin tethering, chromatin contacts with both the lamina and other chromatin segments are largely unaltered by nuclear deformations (**Fig. 4a** and **S13-15**). These measurements show that, in our model, chromatin-lamina tethering allows external mechanical perturbations to be transmitted into peripheral heterochromatin and spatially reorganize the genome.

**FIG. 4.**
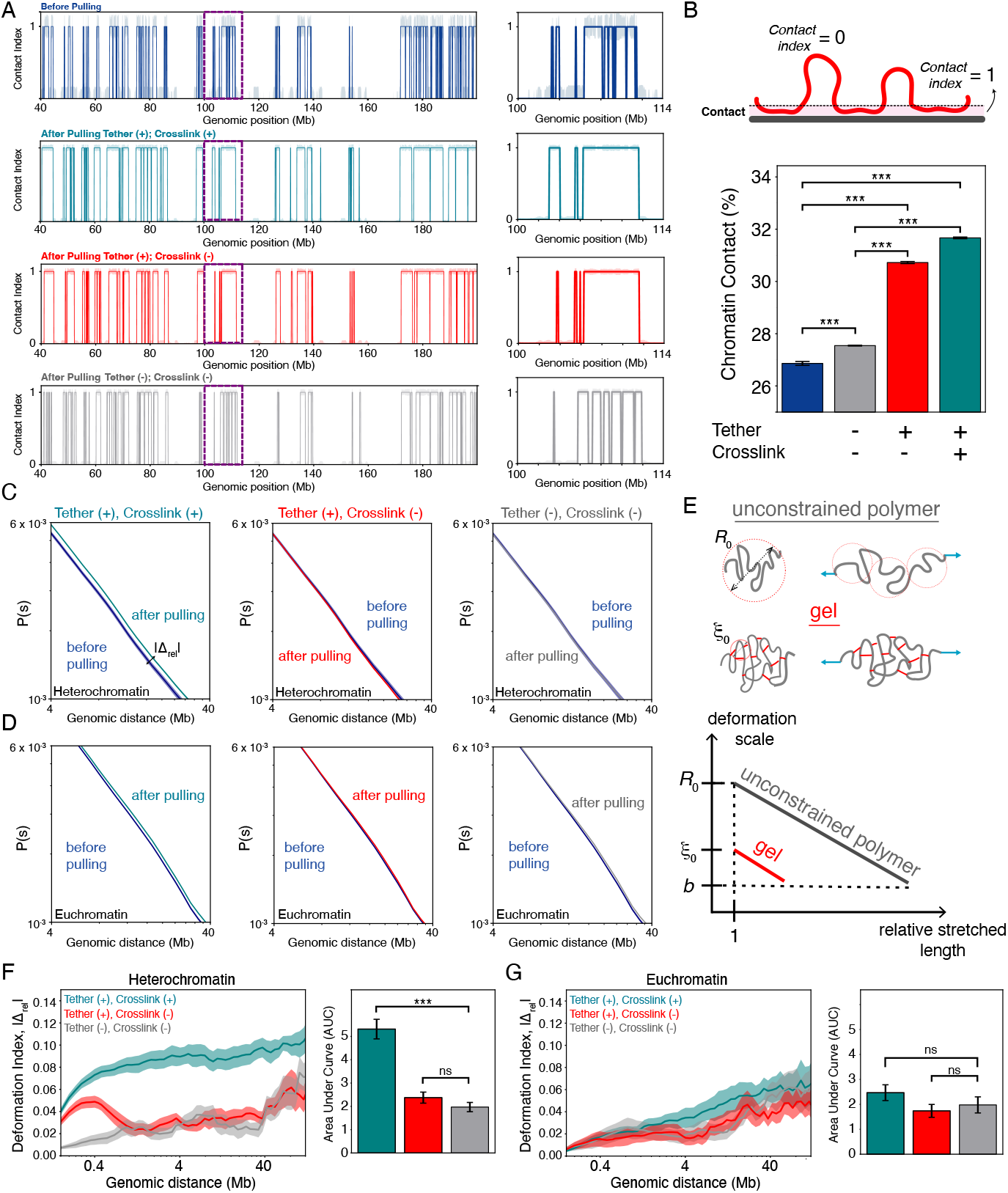
Chromatin-lamina tethering and crosslinking directs chromatin reorganization during nuclear deformation. **A)** Simulated DamID of a 160 Mb region of chromosome 1 showing contact index of genomic sites before stretching (blue) and after 30% axial strain in nuclei with tethering and internal crosslinking of facultative heterochromatin (turquoise), tethering facultative heterochromatin without crosslinking (red), and untethered and uncrosslinked heterochromatin (gray). The contact index is 1 for chromatin subunits in contact with the lamina and 0 otherwise. The solid line shows a representative example, while shading indicates standard error. *Right* : 14 Mb snippet showing zoomed-in view of chromatin-lamina contacts for the region boxed on the left. **B)** *Top*: Illustration of contact index. *Bottom*: Percentage of heterochromatin in contact with the lamina genome-wide, defined as distance *r <* 2.5*σ*, before stretching (navy blue bar) or after pulling to 30% strain in nuclei with and without heterochromatin tethering and/or crosslinking (gray, red, and turquoise).*indicates *p <* 0.05, ** indicates *p <* 0.01, and *** indicates *p <* 0.001. **C)** Chromatin contact frequencies as a function of genomic distance for intra-heterochromatin contacts for the genomic distances of 4-40 Mb, before and after 30% nuclear strain for nuclei with both lamina tethering and heterochromatin crosslinking (left), tethering but no crosslinking (center), or neither tethering nor crosslinking (right). |Δ_rel_| = |(*P*_after_(*s*) − *P*_before_(*s*))*/P*_before_(*s*)| indicates the absolute relative change in *P* (*s*) after deformation. **D)** Chromatin contact frequencies as a function of genomic distance for intra-euchromatin contacts before and after nuclear deformation for different tethering and crosslinking scenarios. **E)** Scaling plot showing the expected size of the internal deformation in chromatin as a function of strain for crosslinked (“gel”) and uncrosslinked (“unconstrained”) polymers. Illustrations show lengthscales over which deformations occur due to presence or lack of constraining linkages. **F)** *Left* : Absolute relative change, |Δ_rel_|, in heterochromatic contacts as a function of genomic distance after deformation of nuclei with both lamina tethering and heterochromatin crosslinking (turquoise), tethering but no crosslinking (red), or neither tethering nor crosslinking (gray). *Right* : Bar plot showing the total absolute relative change, summed over all genomic distances. **G)** Absolute relative change, |Δ_rel_|, in euchromatic contacts as a function of genomic distance, with bar plot showing total change in contacts. Statistical significance is assessed by unpaired, two-sided Student’s t-test.

Remarkably, tethering and crosslinking of peripheral heterochromatin alters how the internal organization of both euchromatin and heterochromatin respond to nuclear deformations. In nuclei with unconstrained chromatin (no tethering and crosslinking), genomic contacts are only minimally perturbed by nuclear deformations of 30% strain (**Fig. 4c-g**). In contrast, with tethering and crosslinking, nuclear deformation alters heterochromatic contacts compared to nuclei with unconstrained chromatin, thus transmitting mechanical signals to chromatin segments. Tethering alone also alters heterochromatic contacts, but to a lesser degree and only for shorter-ranged contacts (*<* 1 Mb). In euchromatin, deformation has a smaller effect on genomic contacts. Nonetheless, increased force transmission with peripheral chromatin crosslinking and tethering leads to a small additional alterations to euchromatic contacts. This difference disappears for larger strains (**Fig. S14**). Together, these observations suggest that regulation of heterochromatin and its connections to the nuclear lamina allow the nuclear interior to sense external forces, while also protecting euchromatin from mechanical disruptions.

## IV. DISCUSSION

### A. Using force to see inside nuclei

Force can be used to biophysically *perturb* cell nuclei, but because nuclei deform in response, force can also *measure* nuclear mechanical properties. Changes in heterochromatin within the nucleus alter nuclear rigidity [3, 26–29, 33, 38, 79]; therefore, forces can be used to *detect* the physical state of the nuclear interior, allowing us to “see” inside the nucleus. Using an established polymer simulation model (**Fig. 1**) [26, 29, 53, 54], we find that physical linkages between peripheral heterochromatin and the nuclear lamina are essential for force transmission into the nucleus and external sensing of the properties of the nuclear interior (**Fig. 3**). This finding complements the biological findings of the importance of peripheral chromatin tethering for mechanotransduction and sequestration and silencing of genes [39, 40, 42, 43, 71, 76].

Our molecular dynamics simulations of nuclei stretched as in micromanipulation experiments [26, 31, 59, 64] reveal the physical mechanism of heterochromatin-based nuclear rigidity. We find that heterochromatin levels, self-affinity, and crosslinking alone cannot regulate nuclear mechanical response (**Fig. 2**). Peripheral heterochromatin *must also* be physically linked to the lamina, so that forces can be elastically transmitted from the lamina to chromatin. In the absence of tethering, simulated nuclei are softer (**Figs. 2-3**). With tethering, heterochromatin crosslinking, but not nonspecific self-affinity, bolsters nuclear stiffness **(Fig. 3)**. Heterochromatin levels are important to the extent that they govern the quantity of chromatin-to-lamina linkages and intra-chromatin crosslinks **Fig. 3**). Consistently, heterochromatin tethering and crosslinking facilitates reorganization of LADs and force transmission to the nuclear interior in response to nuclear deformations (**Fig. 4**). Thus, our model establishes how forces applied to the cell nucleus can be transmitted into the genome to potentially sense and transduce mechanical signals, and how regulation of peripheral heterochromatin might alter mechanobiological response of the nucleus.

### B. Tethering links chromatin to nuclear force transmission and response

Chromatin-lamina tethering profoundly impacts nuclear mechanical response in simulations. In experiments, nuclear mechanics generally exhibits two regimes: chromatin-dominated stiffness in response to small nuclear deformations and lamin-dominated stiffness in response to large deformations [26, 31, 59, 64, 80]. Heterochromatic histone modifications, such as H3K9me3 and H3K27me3, stiffen cell nuclei, while histone hyperacetylation leads to softer nuclei [3, 26–29, 33, 64, 80–82]. Although chromatin modifications alter nuclear stiffness independently of lamins, our results indicate that the physical coupling of chromatin to the lamina is essential for these effects.

Peripheral linkages transmit forces applied to the cell nucleus directly to chromatin to elicit a resistive force response. This could not be achieved if the chromosomes acted as unconstrained polymers. Tethers constrain the polymer chromosomes so that lamina (and nuclear) deformations reposition tethered loci, thus stretching the chromosomal segments between tethered sites (**Fig. 1**). With untethered chromosomes, forces are transmitted globally across the entire chromatin-lamina interface, and unconstrained polymer chromosomes reposition and internally reorganize without significantly stretching chromosomal segments (**Fig. 4**) [72]. This phenomenon has been observed in *Schizosaccharomyces pombe* nuclei depleted of LEM-domain protein orthologs Heh1 and Heh2, which tether chromatin to the nuclear envelope [75]. Nuclei in these cells exhibited larger shape fluctuations and lower-viscosity response to applied forces, consistent with reduced nuclear stiffness and increased chromatin flow. Thus, constraints from chromatin tethering maintain the mechanical integrity and organization of the nuclear periphery by directly transmitting forces to chromatin and enabling elastic response.

Our predictions can be tested by measuring nuclear mechanical response after histone hypermethylation in cells with disrupted chromatin-lamina tethering (*e*.*g*., Sun1/2, emerin, or PRR14 depletion or mutation). In this context, histone hypermethylation (*e*.*g*., via methyl-stat treatment), which normally stiffens nuclei [26–29], should have a smaller effect or even no effect on nuclear rigidity. Similarly, in mouse rod cells, which have an inverted chromatin architecture with heterochromatin in the interior due to loss of tethering by LBR [40, 52], we would expect little change in nuclear stiffness after increasing histone methylation. These experiments could test whether cells can regulate chromatin-lamina tethering as a mechanism for controlling the mechanical functions of heterochromatin.

### C. Peripheral heterochromatin as a tethered polymer gel

An ongoing question is whether chromatin is a polymer fluid or gel [1, 17]. Current evidence suggests that chromatin mechanics is spatially inhomogeneous [30–32], which potentially reconciles evidence for gel-like chromatin [26, 29] with measurements of softer chromatin viscoelasticity [23, 31]. Our simulations suggest that peripheral heterochromatin is tethered to the lamina in a gel-like state and thus can transmit stress across length scales, but the euchromatic interior may be less constrained (*i*.*e*., none and lower levels of crosslinking as compared to heterochromatin).

Heterochromatin tethering is necessary to recapitulate experimental findings that an increase in heterochromatin by histone methylation increases nuclear stiffness [26, 27, 29] (**Figs. 2-3**). Crosslinking of peripheral heterochromatin strengthens this effect (**Fig. 3**). As with chromatin-lamina tethering, crosslinking within heterochromatin spatially constrains chromatin polymer segments so that forces induce stretching and elastic response rather than repositioning by viscous flow. Consistently, chromatin-bridging proteins Cbx5/HP1*α* and BANF1/BAF are necessary for gene stretching and transcription induced by external force [8]. Crosslinks also enable heterochromatin to stiffen cell nuclei independent of further alterations to chromatin-lamina tethering or localization (**Fig. 3**). Similarly, it has been shown that degrading the crosslinker HP1*α* weakens cell nuclei, but does not alter peripheral heterochromatin localization [29]. Thus, spatially uniform crosslinking from previous models [26, 29, 53] is unnecessary; instead, spatially nonuniform, peripheral crosslinking along with tethering is sufficient to drive chromatin-based nuclear mechanical response.

Uncrosslinked interior euchromatin (**Fig. 1**), in contrast, might allow the relatively soft elastic and viscous response observed in experiments probing the chromatin interior on shorter lengthscales [23, 24]. Spatial inhomogeneity in chromatin material properties is maintained by histone modifications and other genomically and spatially inhomogeneous patterning [42, 71, 83, 84]. Interior euchromatin exhibits histone acetylation, which leads to softer elastic response in force spectroscopy studies [26, 27, 64] and liquid-like properties for *in vitro* chromatin [18]. However, a simple uncrosslinked polymer model for euchromatin remains at odds with other observations of chromatin dynamics. Rheological measurements with magnetic beads suggest crosslinking on lengthscales of half a micron [25]. This may suggest an intra-chromatin crosslinking with a relatively large crosslinking length, but observations of loss of structure in “liquid chromatin” Hi-C experiments suggest linkages every few kb [85]. A model with dynamic crosslinking (a “weak gel”) [23, 72, 86–88] may better explain chromatin’s viscoelastic properties, but remains to be extensively explored in the context of chromatin mechanics.

### D. Force transmission through chromatin tethers and nuclear function

Force transmission through chromatin-lamina linkages is important for proper mechanotransduction of physical stresses into biochemical processes [1, 43]. Experiments using magnetic beads to apply forces to mammalian cells have shown that external forces can be transmitted to chromatin, stretch individual genes, and elevate gene expression [8, 89]. These effects depend on chromatin-lamina linkages, such as SUN1/2 and emerin. Our polymer simulations show that chromatin tethers are necessary for force transmission and activation of chromatin’s elastic response (**Figs. 3** and **4**). This hints that nuclear mechanical properties could potentially guide the transduction of mechanical cues into biochemical and gene regulatory activity.

Remarkably, our analyses revealed that tethering to the lamina leads to large deformations of heterochromatin during nuclear strain. We observe this effect despite comparatively small alterations in euchromatin organization (**Fig. 4**). This defied our initial expectation that applied forces would induce larger deformations and changes in chromatin contacts in the more flexible euchromatin than in the stiffer heterochromatin. However, the simulations are consistent with experimental observations of genomes altered by nuclear deformations. In Hi-C experiments with cells that migrated through pores smaller than the nuclear diameter (5 *µ*m), a few percent of all genomic regions switched between euchromatic A- and heterochromatic B-type compartments, as defined by chromatin contacts [5, 6]. Switches from B to A were observed more frequently, possibly indicating significant changes in B, and the relative frequency of chromatin contacts within B compartments was lower after migration. Similarly, in simulations of nuclei deformed by axial tension, peripheral heterochromatin tethering combined with heterochromatin crosslinking favored changes in heterochromatin contact frequencies at ~ 1 Mb scales (**Fig. 4**). These alterations occurred because tethering and crosslinking facilitated force transmission through heterochromatin. These findings are consistent with deformation microscopy imaging showing that denser chromatin and peripheral H3K9me3-marked heterochromatic regions undergo larger strains than lower density, transcriptionally active euchromatic regions during cardiomyocyte contraction [15, 30]. We conclude that physical linkages not only stiffen nuclei but also redistribute strain.

Consequently, regulation of heterochromatin through both crosslinking and peripheral heterochromatin may be essential to mechanobiological responses to external forces. Our observations of force transmission through the nucleus suggest that polymer physical constraints, such as heterochromatin tethering and crosslinking (**Fig. 4**), could be mechanistically important to nuclear mechanosensing [1, 8, 43, 89]. Complementarily, tethering is necessary to detect the internal chromatin architecture via external mechanical perturbations (**Fig. 3**). This tethering promotes heterochromatin-based nuclear rigidity, which can protect the genome from damage [28]. However, the corresponding increase force transmission can *increase* DNA damage [3]. These competing effects may depend on the composition (*e*.*g*., lamin A level) and overall rigidity of the nucleus [2, 3], as well as the scale of deformation (**Figs. 4** and **S12**). Our model suggests that cells may tune these competing mechanical effects through regulation of heterochromatin quantities, crosslinking, and tethering to the lamina.

## CODE AVAILABILITY

The codes used for this study are freely and publicly available at https://github.com/agattar/Attar_Mechanics_Paper.

## ACKNOWLEDGEMENTS

We thank Andrew Stephens for fruitful discussions. EJB acknowledges support from the NIH Common Fund 4D Nucleome Program (UM1HG011536). This work has been supported by The Council of Science and Technology of Turkey (TUBITAK) Grant no 122F309 and the National Science Center, EU’s H2020 Programme and MSCA Grant Agreement no. 945339 Poland [Grant Polonez Bis No. 2021/43/P/ST3/01833]. The numerical calculations reported in this paper were fully performed at TUBITAK ULAKBIM, High Performance and Grid Computing Center (TRUBA resources).

## AUTHOR CONTRIBUTIONS

All authors participated in writing and reviewing the manuscript.

