## Supplementary_File for "Peripheral heterochromatin tethering is required for chromatin-based nuclear mechanical response"

#### The Molecular Dynamics Simulation Parameters

For modeling the non-bonded pairwise interaction between the beads, we employed Lennard-Jones (LJ) potential. LJ potential was shifted and truncated as in the following,

$$V(r) = u\left[\left(\frac{\sigma}{r}\right)^9 - \left(\frac{\sigma}{r}\right)^6 + v_s\right], \quad r \le r_c \quad V(r) = 0, \quad r > r_c$$

The cutoff distance was set to $r_c = 2.5\sigma$ which $\sigma$ corresponds to the unit length in our model, and the shift factor in the potential was set to $v_s = 0$. The attraction strengths used in the simulations follow the definitions from our previous work [1] and are listed in Table 1.

The interactions in the model are repulsive with a repulsive interactions strength of $u = 1kT$, a cutoff distance of $r_c = 2^{\frac{1}{6}}\sigma$ and a shift factor of $v_s = \frac{1}{4}$. Unless specified otherwise, the timestep defined for the simulations was set as $\Delta t = 0.005\tau$, $\tau$ corresponds to the time unit. Brownian dynamics were implemented through the Langevin thermostat in our simulations with the microcanonical ensemble (*NVE),* and ensured constant temperature. The damping coefficient for Langevin thermostat was set to 1000τ unless stated otherwise. Periodic boundaries were applied in all directions, and the simulation box dimensions were defined as $60\times60\times60\sigma^3$. All simulations were run using the LAMMPS MD package.

For the bonded interactions, two different approaches were employed. Except for the bonds used for crosslinking within constitutive and facultative heterochromatin beads, all other assigned bonds followed the non-extensible FENE potential.

$$V(r) = -0.5kR_0^2 ln[1 - \left(\frac{r}{R_0}\right)^2]$$

The maximum bond distance, maximum extension, for the FENE potential was defined as $R_0$ =1.5. The bond energy was set to $k = 5\frac{kT}{\sigma^2}$ for shell beads while for chromatin polymer, bond energy was set to $k = 30\frac{kT}{\sigma^2}$. For crosslinking of the constitutive and facultative heterochromatin bead, harmonic bonds were used as described in the following,

$$V(r) = k(r - r_0)^2$$

where the bond distance set as $r_0$ = 1.5σ and bond energy was set to $k = 10\frac{kT}{\sigma^2}$. For the molecular tethers assigned for tethering the heterochromatin domains, the energy of the bond was set to $k = 10\frac{kT}{\sigma^2}$.

Polymer flexibility was achieved through defining the harmonic angle potential. Since the heterochromatin posseses longer persistence lengths (100–200 nm) than that of euchromatin (10–50 nm), angle restrictions were defined to facultative heterochromatin beads which the potential defined as in the following,

$$V(\theta) = k_\theta(\theta - \theta_0)^2$$

where the energy, $k_\theta$, was defined as $k_\theta = 1\frac{kT}{rad^2}$ and the angle was set to $\theta = 180$ to ensure a semiflexible heterochromatin.

| Type | Interaction Energy |
|---|---|
| Euchromatin - Euchromatin | 0.05 kT |
| Euchromatin - Facultative Heterochromatin | 0.25 kT |
| Euchromatin – Cons. Heterochromatin | 0.50 kT |
| Fac. Heterochromatin – Fac. Heterochromatin | 0.65 kT |
| Heterochromatin-Shell | 0.75 kT |
| Fac. Heterochromatin – Cons. Heterochromatin | 0.85 kT |
| Cons. Heterochromatin – Cons. Heterochromatin | 1.10 kT |
| Telomere-Shell | 2.20 kT |

**Table 1** The interaction energies set for various beads in the simulations where k and T are Boltzmann and absolute temperature.

#### System Design

The initial model is designed following on our previous work [1]. The 10 different configurations, with various initial placement of each chromosome providing the randomness in our simulations, in the final equilibration step of the simulated conventional nucleus simulations were used.

For the tether assignments, only the beads within the contact of the lamina, $\sigma \leq 2.5$, were considered. Each bead was provided with a capacity of only a single tether to the random lamina bead within the proximity.

For crosslink assignments, beads within a contact proximity, $\sigma \leq 2.5$, were considered. The probability factor, $p_C$, was set to 0.2, unless otherwise indicated, which each bead had a 20% chance of being chosen for crosslinking. The chosen beads were assigned with a minimum of one and maximum of two crosslinks to another, randomly selected, bead within the contact proximity. Crosslinks assignments were only allowed within the same chromatin type, for instance only between facultative heterochromatin domains.

For perturbation conditions, to decrease the facultative heterochromatin fractions, facultative heterochromatin beads were randomly selected and converted to the euchromatin bead type, achieving a final $f \approx 20\%$ facultative heterochromatin fraction. On the other hand, to increase the facultative heterochromatin fractions, euchromatin beads that were positioned below $10\sigma$ of average radial distance of nuclear lamina were selected and converted to the facultative heterochromatin bead type, achieving a final $f \approx 60\%$ facultative heterochromatin fraction.

#### Analysis of Simulation Trajectories

##### Nuclear Spring Constant Calculation

The external force applied to the system on the y-axis by *fix smd* was extracted through *fix ave/time* command. The strain was calculated by the maximally displaced lamina bead respectively to the origin. For nuclear spring constant calculations, 5 different strains were selected from 10% to 50%. For calculating of the nuclear spring constants, the local slope was calculated by the following:

$$k_{nuc} = \frac{F_{\epsilon} - F_{\epsilon-0.1}}{\epsilon - (\epsilon - 0.1)} = \frac{F_{\epsilon} - F_{\epsilon-0.1}}{0.1}$$

**Simulation of a DamID Map**

To calculate the heterochromatin-lamina contacts, the same distance proximity, $\sigma \leq 2.5$, was used. The beads having contacts with lamina beads are assigned as a contact by arbitrary contact index, 1. Simulated DamID maps were generated at 30% strain using 10 replicates and 11 configurations for each replicate, generating 110 configurations in total. Configurations are selected by including the 5 next and previous timesteps from the 30% strain timepoint.

**The Calculation of the Contact Frequency as a Function of Genomic Distance, P(s)**

To analyze chromatin contacts across varying genomic distances, we computed the probability of spatial proximity for different genomic separations, s. For that purpose, the constitutive heterochromatin domains were not included and each polymer chain, remaining with $N$=5000 (discarding the telomere beads), was processed separately for calculations, and then summed together. We defined a series of genomic distances, s, starting from s=4, using a geometric progression with an approximate growth factor of 1.12 resulting in a sequence of [4, 5, 6, 7, 8, 9, 10, 11,…., 1833, 2049, 2290, 2560, 2862, 3199, 3576, 3998, 4469, 4996 (N-4)] [2]. For each genomic distance, the number of contacts, satisfying the distance threshold $\sigma \leq$ 2.5, were calculated for each bead in the chain and divided by the genomic distance, s. The resulting contact probability matrix is then normalized by the total chain length, except constitutive heterochromatin, in our model. Additionally, the same approach was followed for calculating the P(s), for different chromatin types. For that case, chromatin contacts were calculated separately both for euchromatin and facultative heterochromatin, considering only the intra-contacts for each chromatin type. Then, the contact probability vector was normalized separately, divided by the total number of facultative heterochromatin or euchromatin. These were generated at 30% strain using 10 replicates and 11 configurations for each replicate, generating 110 configurations in total, unless stated otherwise. Configurations are selected by including the 5 next and previous timesteps from the 30% strain timepoint.

#### Appendix

Subsections of a chain have ideal chain statistics at length scales below the thermal length ($\xi_T$), and above the correlation length ($\xi_C$). [3] The ideal chain statistics at these length scales is due to screening of two-body interactions (whether attractive or repulsive) by thermal fluctuations at length scales below the thermal length, and by concentration fluctuations at length scales above the correlation length. While thermal length is determined by the temperature, correlation length is controlled by the concentration. Correlation length (mesh size) decreases with increasing concentration, and becomes smaller than thermal length in the concentrated regime of $\phi > \phi''$. Thus, chains in a concentrated solution have ideal chain statistics at all length scales (see **Figure 1**).

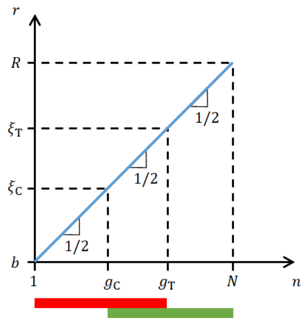


**Figure 1:** Log-log plot of root-mean-square end-to-end distance $r(n)$ of a chain section of $n$ Kuhn monomers of size $b$ each. The $n$-mer sections in the red region (sections smaller than the thermal length $\xi_T$) and in the green region (sections larger than the correlation length $\xi_C$)

have ideal random walk conformations of $r(n) \simeq bn^{1/2}$. Since $\xi_C < \xi_T$ in a concentrated solution, the chain have ideal conformations at all length scales.

##### Stretching a single chain in a concentrated solution of chains

An unperturbed ideal chain of $N$ Kuhn monomers of size $b$ each, has a root-mean-square end-to-end distance of $R_0 \simeq bN^{1/2}$. The external work done to stretch the chain is not spent to decrease the entropy of the chain at all length scales by aligning all the bonds a bit in the stretching direction, but instead the ideal chain conformations are perturbed only above a certain length scale, known as the tension length, or size of a Pincus blob ($\xi_P$). [4] The chain sections inside a Pincus blob of size $\xi_P$ are still ideal random walks, but the whole chain is an extended array of such Pincus blobs (see **Figure 2**).

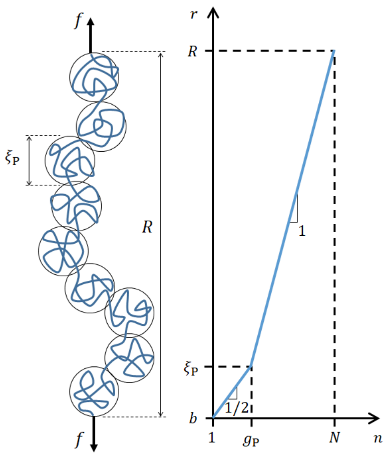


**Figure 2:** A stretched chain is an extended array of Pincus blobs of size $\xi_P$ each. Size $r(n)$ of an $n$-mer subsection of the chain is shown in a log-log plot.

As the stretching force gets larger, smaller sections of the chain get perturbed, i.e., the number of monomers in a Pincus blob ($g_P$) decreases, and hence, Pincus blobs get smaller. But the number of Pincus blobs per chain ($N/g_P$) increases, and as a result, chain size increases.

The force $f \simeq kT/\xi_p$ required to bring the size of the chain from $R_0$ to $R = \alpha R_0$ can be calculated as [3]

$$f \simeq \kappa_{ch} \alpha, \tag{1}$$

where $\alpha = R/R_0$ is the elongation ratio and $\kappa_{ch} \simeq kT/R_0$ is the chain stiffness.

**Stretching a gel in the concentrated regime**

Consider we crosslink the ideal chains in a concentrated solution to form a polymer network. Let $\xi_0$ denote the average distance between network strands (correlation length, mesh size) in the absence of external forces. Even though no external force is acting on the gel at equilibrium, crosslinking the chains introduces a tensile stress on network strands that is on the order of $kT/\xi_0^3$. This elastic stress tries to collapse the gel. On the other hand, an osmotic pressure $kT/\xi_0^3$, due to three-body interactions, tries to expand the gel. At equilibrium in the absence of external forces, these elastic and osmotic pressures are at a balance. Hence, an undeformed gel has Pincus blobs with the same size $\xi_0$ as the network mesh size. These Pincus blobs shrink further upon deformation of the gel under external forces. Scaling theory predicts a uniaxial force-deformation relation as [5]

$$f \simeq \kappa_{nw} \alpha, \tag{2}$$

where $\kappa_{nw} \simeq kT/\xi_0$ is the gel network stiffness [6].

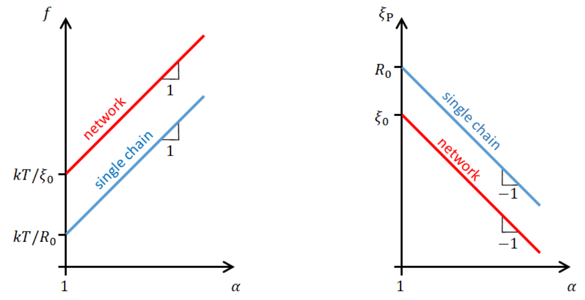


**Figure 3:** Log-log plot of force-elongation relations (left panel) and tensile length-elongation

relations (right panel) for a single chain and for a network of chains.

Although the force-elongation relations for a single chain and for a gel [Eqs. (1) and (2)] are both linear, we have different stiffness expressions for the chain and the gel. Since $R_0$ is larger than $\xi_0$, a single chain ($\kappa_{ch} \simeq kT/R_0$) is softer than a gel network ($\kappa_{nw} \simeq kT/\xi_0$), see **Figure 3**. In both cases, length scales above which the conformations are perturbed (i.e., size of Pincus blobs) decrease with increasing elongation. Combining the relations $\xi_P \simeq b g_P^{1/2}$, $R = \xi_P \left( N/g_P \right)$ and $R = \alpha R_0$, we obtain the size of Pincus blobs as a function of elongation ratio as [3]

$$\xi_P \simeq R_0 \alpha^{-1} \quad (3)$$

for a single chain and

$$\xi_P \simeq \xi_0 \alpha^{-1} \quad (4)$$

for a network of chains. The difference is that, for small deformations $\alpha \approx 1$, Pincus blobs begin from the size $R_0$ for a single chain, and from $\xi_0$ for a network, see **Figure 3**. This difference originates from the tensile stress on network strands, imposed by crosslinking and balanced by the osmotic pressure. These pre-stressed network stands already have a Pincus blob size of $\xi_0$ without any externally applied force.

#### Supplementary Figures

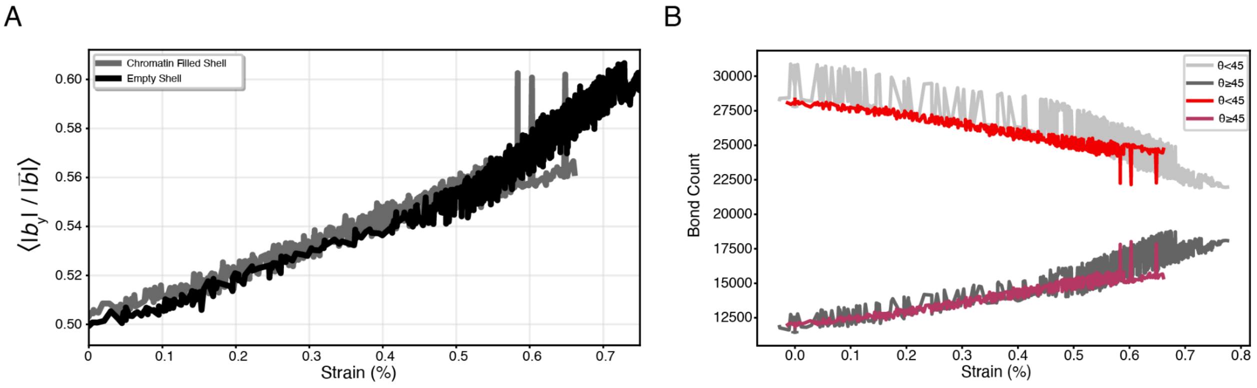


**Figure 4** Nuclear lamina deformation under force. A) Alignment of the lamina bonds across the force direction in empty shells and chromatin filled shells. B) Number of bonds alignment across the force direction.

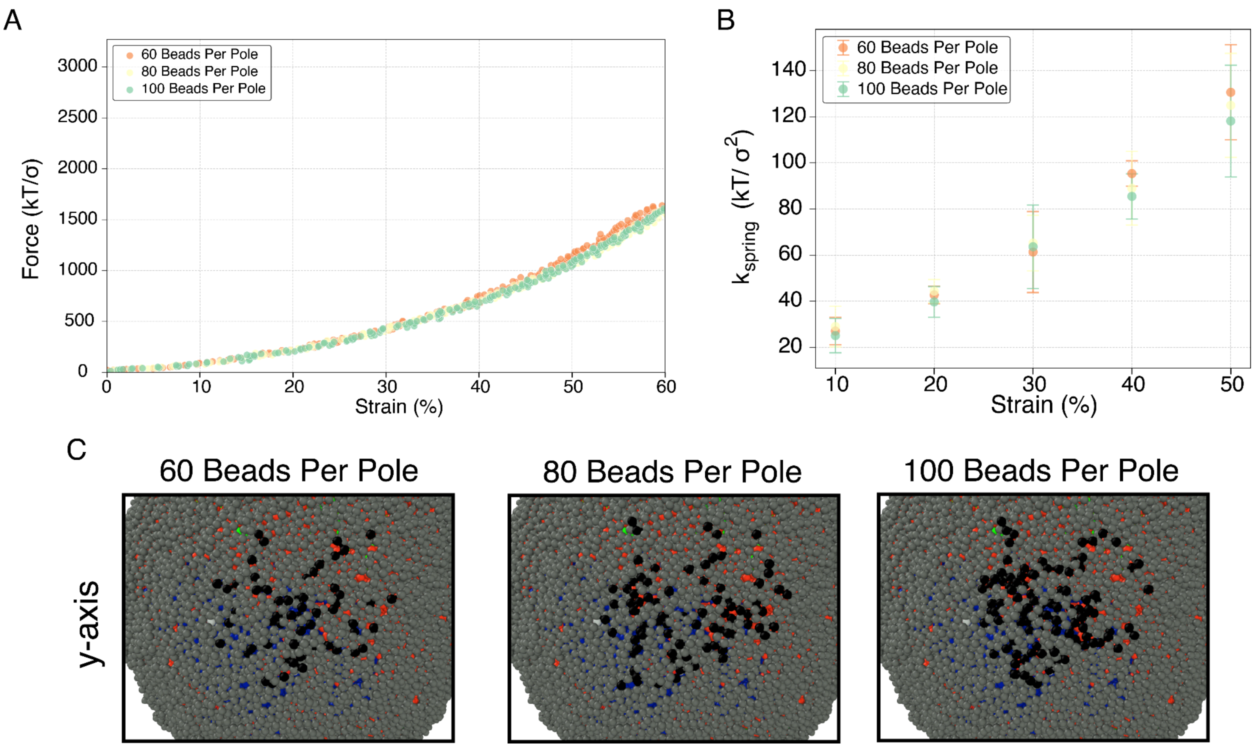


**Figure 5:** Simulation benchmark. A) Force-strain curves of single replicates with varying numbers of beads assigned from the lamina shell for pulling force application. B) Spring constants extracted at 30% strain under various conditions. C) Snapshots of force-exposed beads before shell pulling. n = 3.

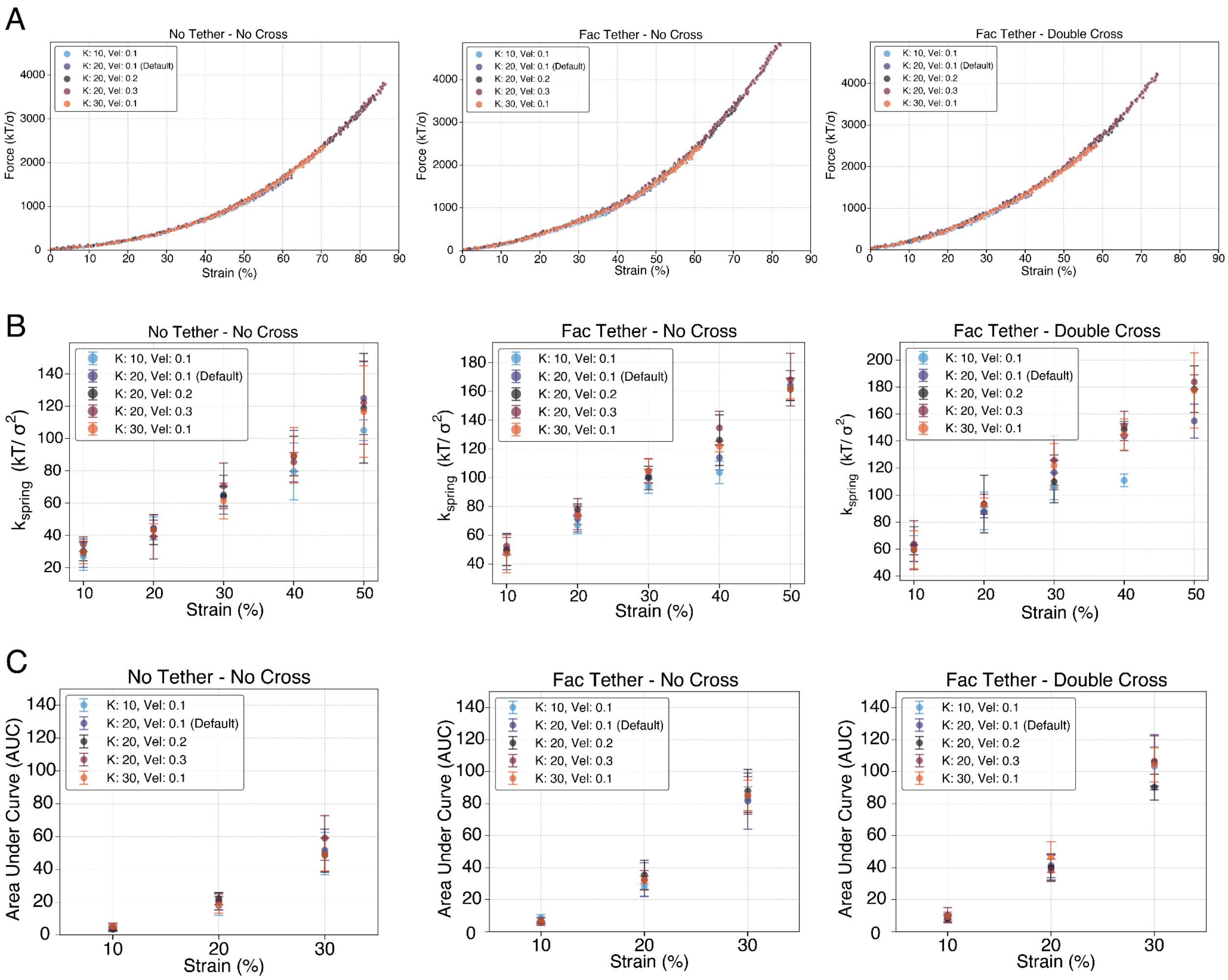


**Figure 6:** Simulation benchmark. A) Force-strain curves of single replicates under different tethering and crosslinking conditions, with varying pulling parameters: spring constant (K) and pulling velocity (vel). B) Spring constants extracted at 30% strain under various conditions and pulling parameters. C) Areas under curves extracted at 30% strain under various conditions and pulling parameters. n=3.

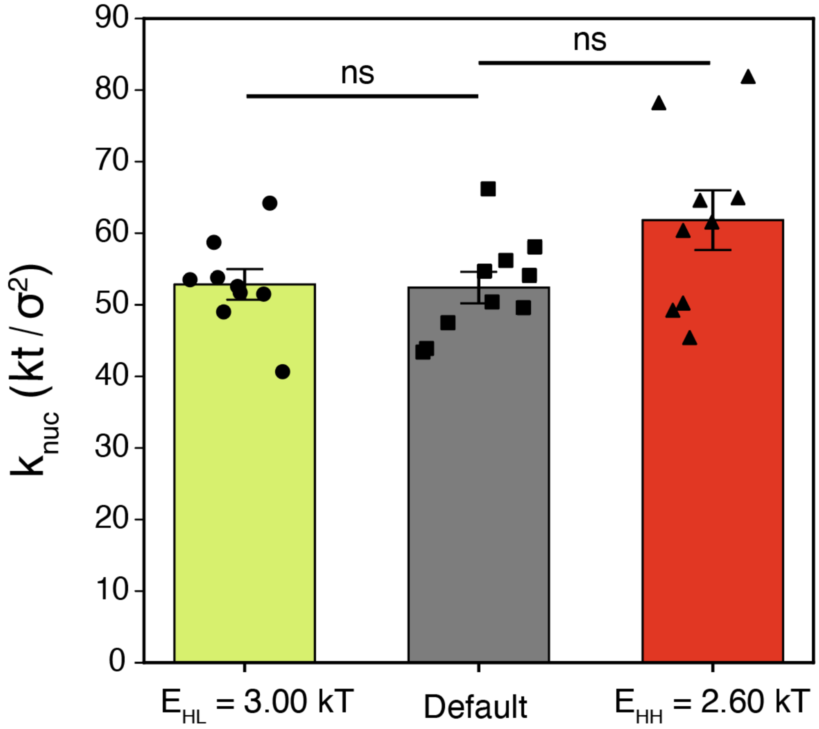


**Figure 7**. Nuclear spring constants for extreme heterochromatin-lamina and heterochromatin heterochromatin interactions at 60% facultative heterochromatin fractions. Bars are mean ± SEM. Statistical significance is assessed by one-way ANOVA followed by a post-hoc Tukey's HSD test for pairwise comparisons. * denotes $p < 0.05$, ** denotes $p < 0.01$, and *** denotes $p < 0.001$. n=10.

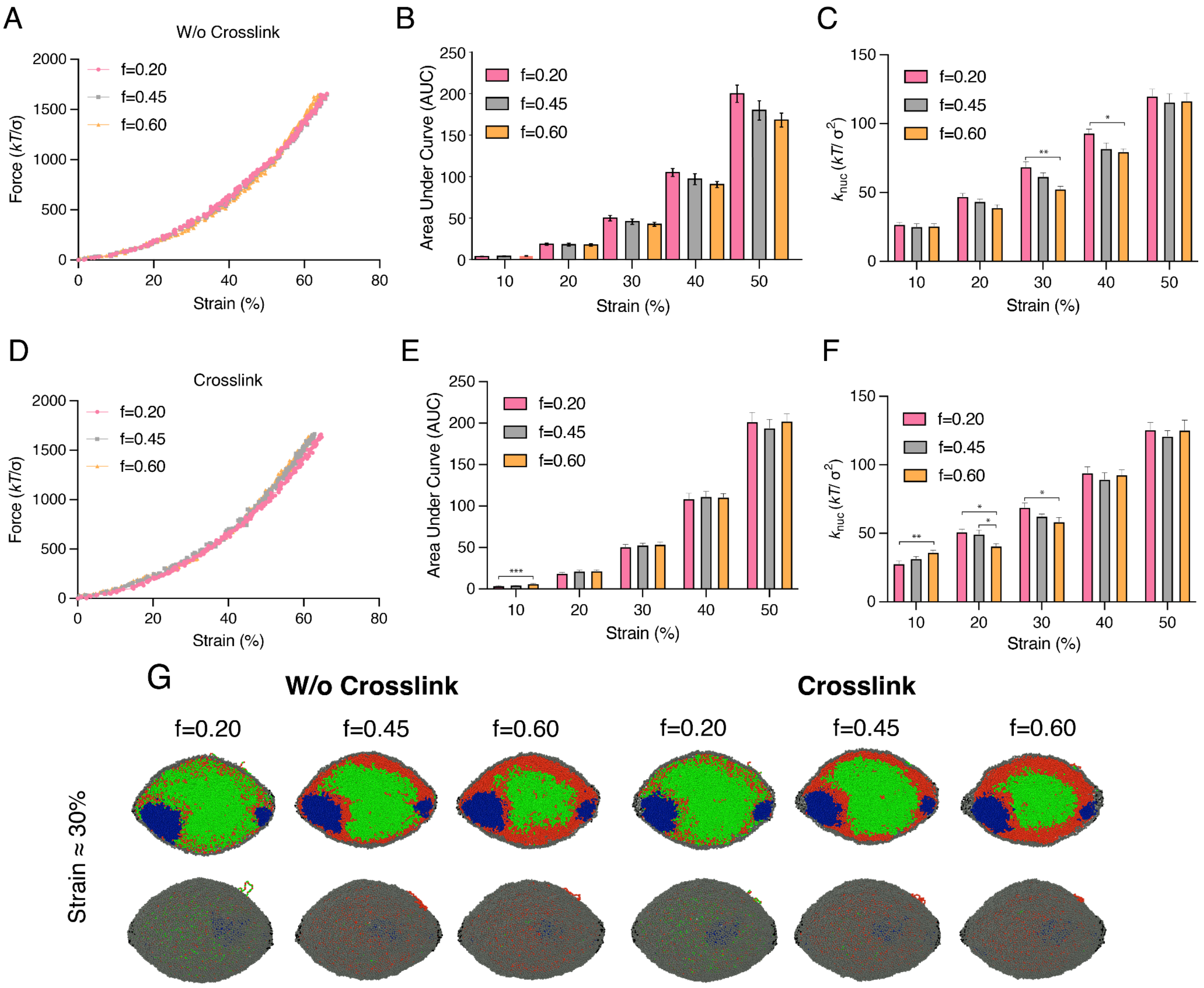


**Figure 8**: Comparison between non-crosslinked and crosslinked heterochromatin domains. A) Force-strain curves of single replicates of various heterochromatin fraction conditions without crosslinking. B) Area under curves extracted from force-strain curves of various facultative heterochromatin fraction conditions without crosslinking at different strain values. C) Spring constant values from different strain values for various facultative heterochromatin fractions without crosslinking. D) Force-strain curves of single replicates of various heterochromatin fraction conditions with crosslinking. E) Area under curves extracted from force-strain curves of various facultative heterochromatin fraction conditions with crosslinking at different strain values. F) Spring constant values from different strain values for various facultative heterochromatin fractions with crosslinking. G) Snapshots showing internal organization and nuclear shell morphology at 30% strain for conditions with and without crosslinking conditions. Bars are mean ± SEM. Statistical significance is assessed by one-way ANOVA followed by a post-hoc Tukey's HSD test for pairwise comparisons. * denotes $p < 0.05$, ** denotes $p < 0.01$, and *** denotes $p < 0.001$. n=10

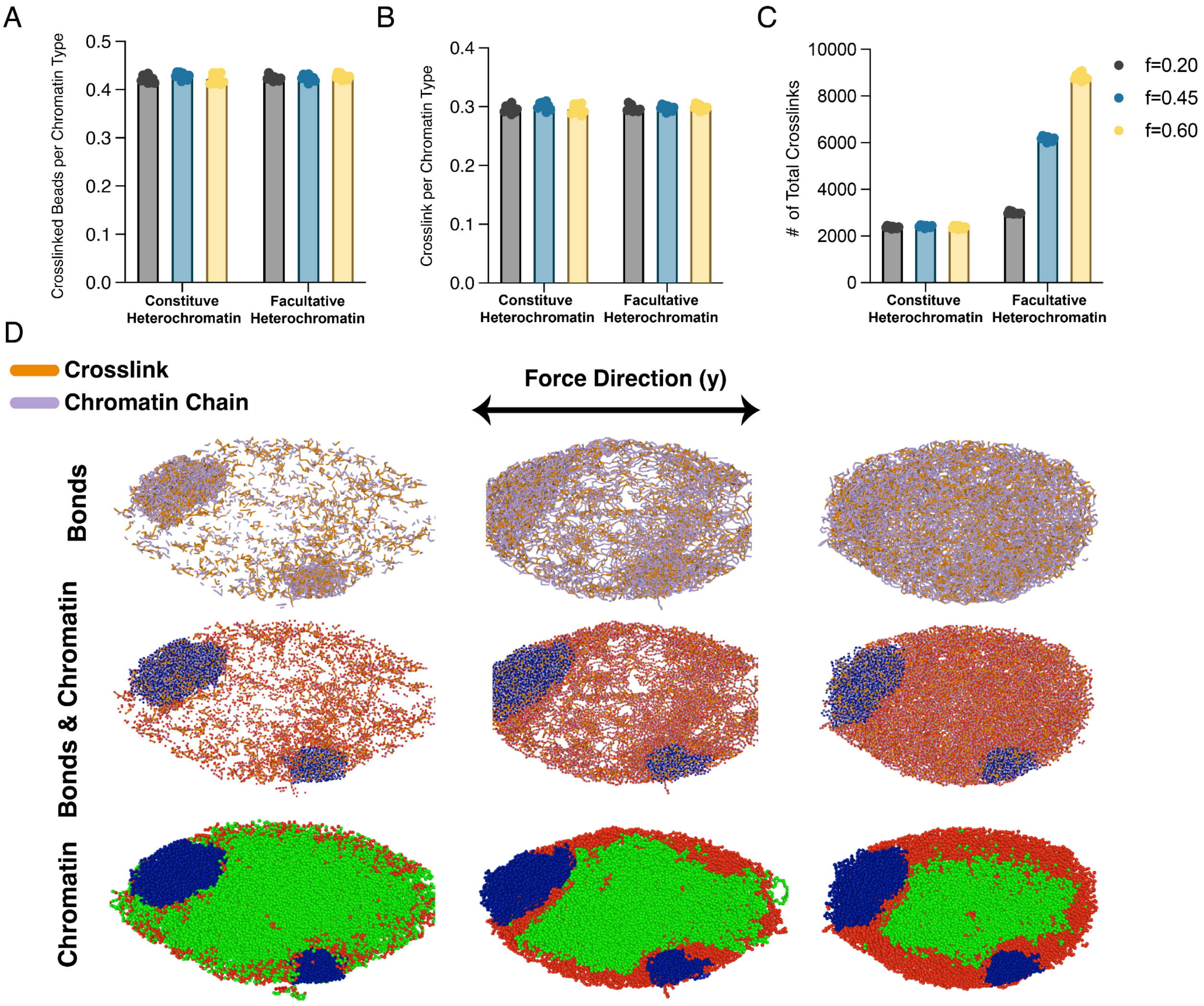


**Figure 9**. Crosslink analysis. A) Bar graphs showing the percentage of constitutive and facultative heterochromatin with crosslinking across different facultative heterochromatin fractions. B) Bar graphs showing the number of crosslinks per each heterochromatin domain. C) Total number of crosslinks assigned to each facultative heterochromatin fraction condition. D) Snapshots of bond and chromatin networks for different facultative heterochromatin fraction conditions at 30% strain. n = 10.

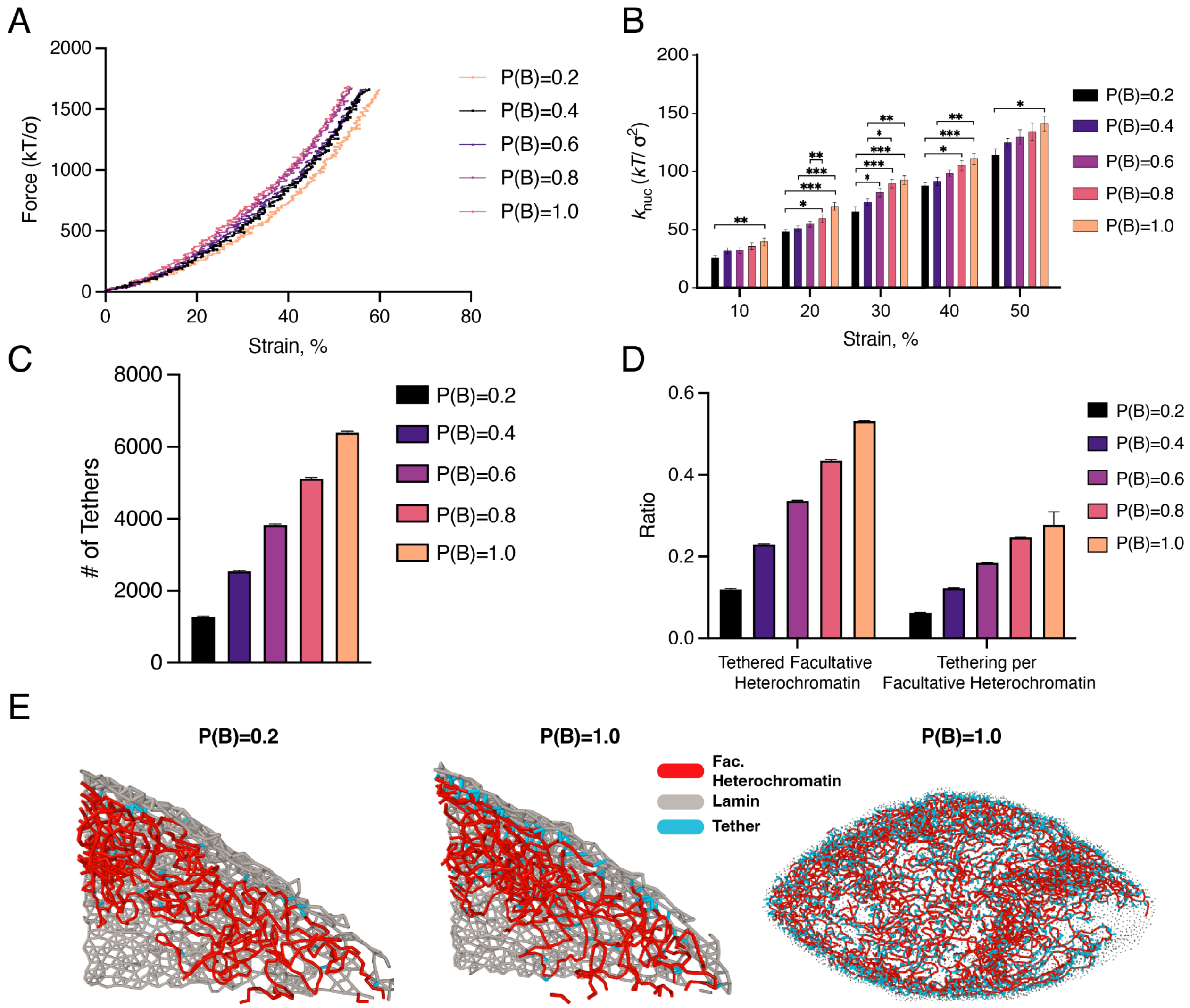


**Figure 10.** Optimization of tether assignment. A) Single replicate force-strain curves for each tethering probability assignment. B) Spring constants at various strain values for different tethering probability conditions. C) Total number of tethers assigned for each probability condition. D) Ratios of tethered facultative heterochromatin and tethering per each facultative heterochromatin are demonstrated by barplot. E) Snapshots showing peripheral facultative heterochromatin organization (red), and assigned tethers (blue). Bars are mean ± SEM. Statistical significance is assessed by one-way ANOVA followed by a post-hoc Tukey's HSD test for pairwise comparisons. $^{*}$ denotes $p < 0.05$, $^{**}$ denotes $p < 0.01$, and $^{***}$ denotes $p < 0.001$. n=10.

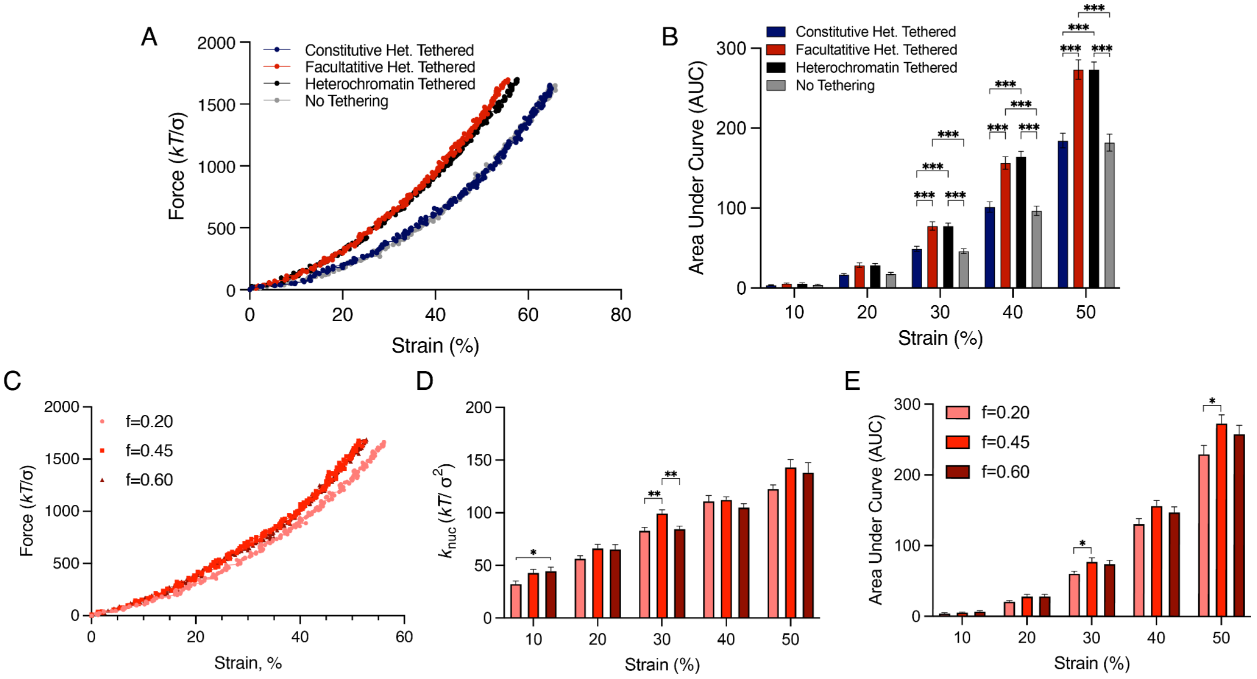


**Figure 11.** Tethering without crosslinking. A) Force-strain curves of single replicates for each tethering condition without crosslinking. B) Area under curves extracted from force-strain curves for different conditions at various strain values. C) A) Force-strain curves of single replicates for various facultative heterochromatin fractions. D) Spring constants for different facultative heterochromatin fraction conditions at various strain values. E) Area under curves extracted from force-strain curves for different conditions at various strain values. Bars are mean ± SEM. Statistical significance is assessed by one-way ANOVA followed by a post-hoc Tukey's HSD test for pairwise comparisons. * denotes $p < 0.05$, ** denotes $p < 0.01$, and *** denotes $p < 0.001$. n=10.

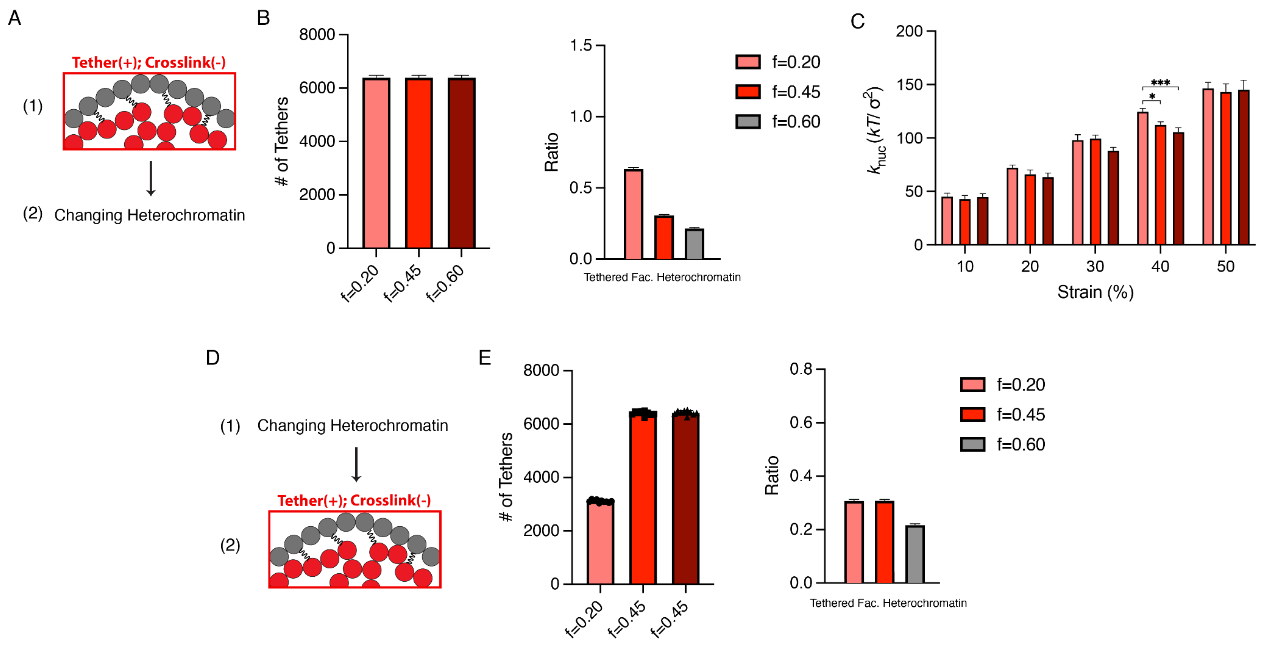


**Figure 12.** Two approaches for perturbation modeling. A) Schematic of the first perturbation approach; tethers and crosslinks are assigned after the final equilibration step, followed by modification of facultative heterochromatin fractions. B) Total number of tethers across different heterochromatin fractions and bar plot showing the ratio of tethered facultative heterochromatin domains. C) Spring constants calculated at various strain values for different facultative heterochromatin fractions. D) Schematic of the second perturbation approach; the facultative heterochromatin fractions are modified after the final equilibrium step, followed by introduction of crosslinks and tethers to the system. E) Total number of tethers for different heterochromatin fractions and bar plot showing the ratio of the tethered facultative heterochromatin domains. Bars are mean ± SEM. Statistical significance is assessed by one-way ANOVA followed by a post-hoc Tukey's HSD test for pairwise comparisons. * denotes $p < 0.05$, ** denotes $p < 0.01$, and *** denotes $p < 0.001$. n=10.

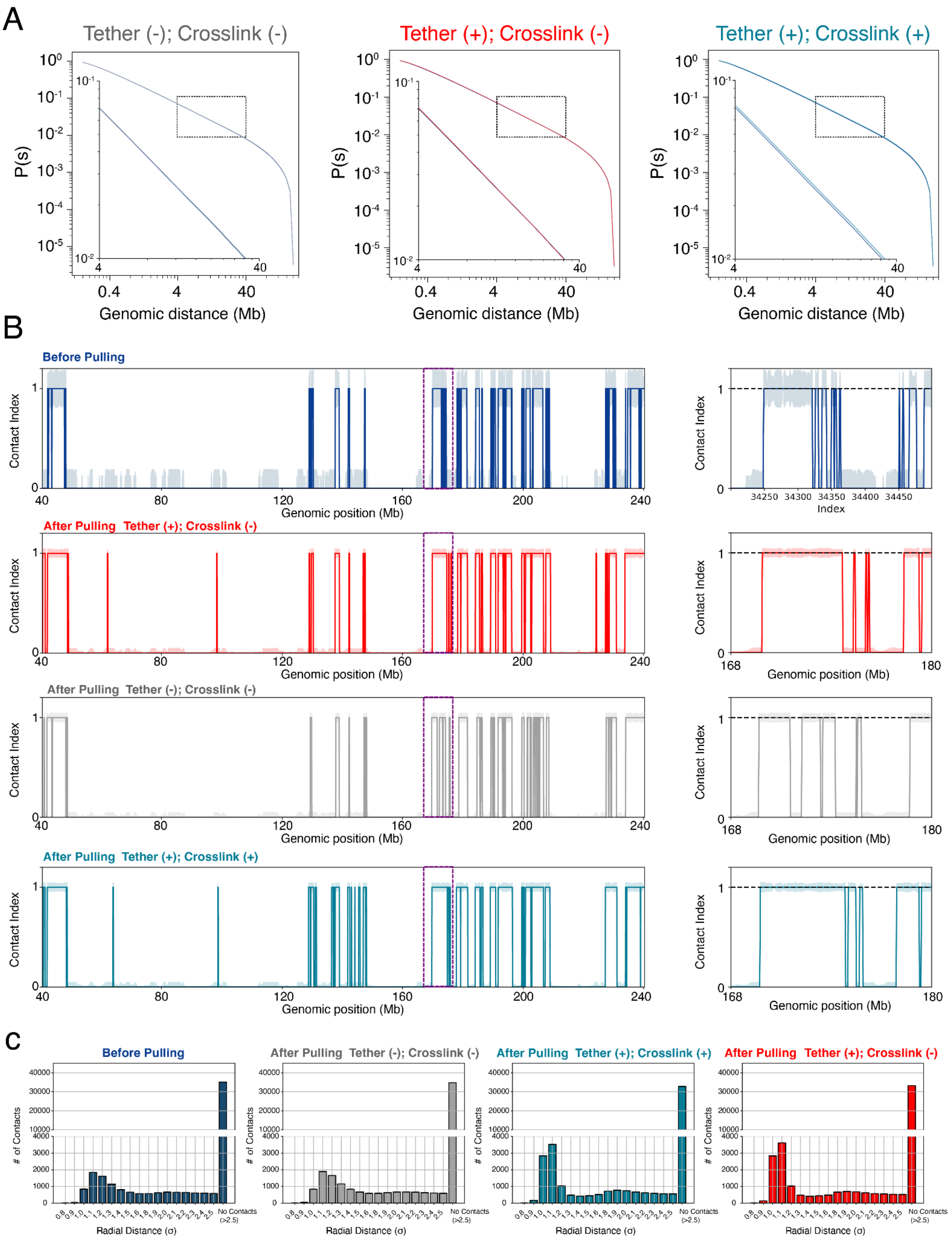
A
Tether (-); Crosslink (-)
Tether (+); Crosslink (-)
Tether (+); Crosslink (+)
P(s)
Genomic distance (Mb)
B
Before Pulling
After Pulling Tether (+); Crosslink (-)
After Pulling Tether (-); Crosslink (-)
After Pulling Tether (+); Crosslink (+)
Contact Index
Genomic position (Mb)
Index
C
Before Pulling
After Pulling Tether (-); Crosslink (-)
After Pulling Tether (+); Crosslink (+)
After Pulling Tether (+); Crosslink (-)
### of Contacts
Radial Distance (σ)
No Contacts (>2.5)

**Figure 13.** Chromatin reorganization during nuclear deformation is regulated by peripheral facultative heterochromatin-lamina tethering and crosslinking. A) Chromatin contact frequencies as a function of genomic distance for intra-facultative heterochromatin contacts for the genomic distances of 4-40 Mb, before and after 30% nuclear strain for nuclei with both lamina tethering and heterochromatin crosslinking (right), tethering but no crosslinking (center), or neither tethering nor crosslinking (left). B) Simulated DamID map for 200 Mb region of the chromosome 2 at 30% strain for before pulling (first row), for nuclei with tethering but no crosslinking (second row), neither tethering nor crosslinking (third row), and both lamina tethering and heterochromatin crosslinking (fourth row). Purple dashed windows represent the genomic position of the snippets exhibiting a contact map for 12 Mb region for each condition. C) The distribution of the number of contacts and the corresponding distance intervals at 30% strain for before pulling (first row), neither tethering nor crosslinking (second), both lamina tethering and heterochromatin crosslinking (third row), and for nuclei with tethering but no crosslinking (fourth row).

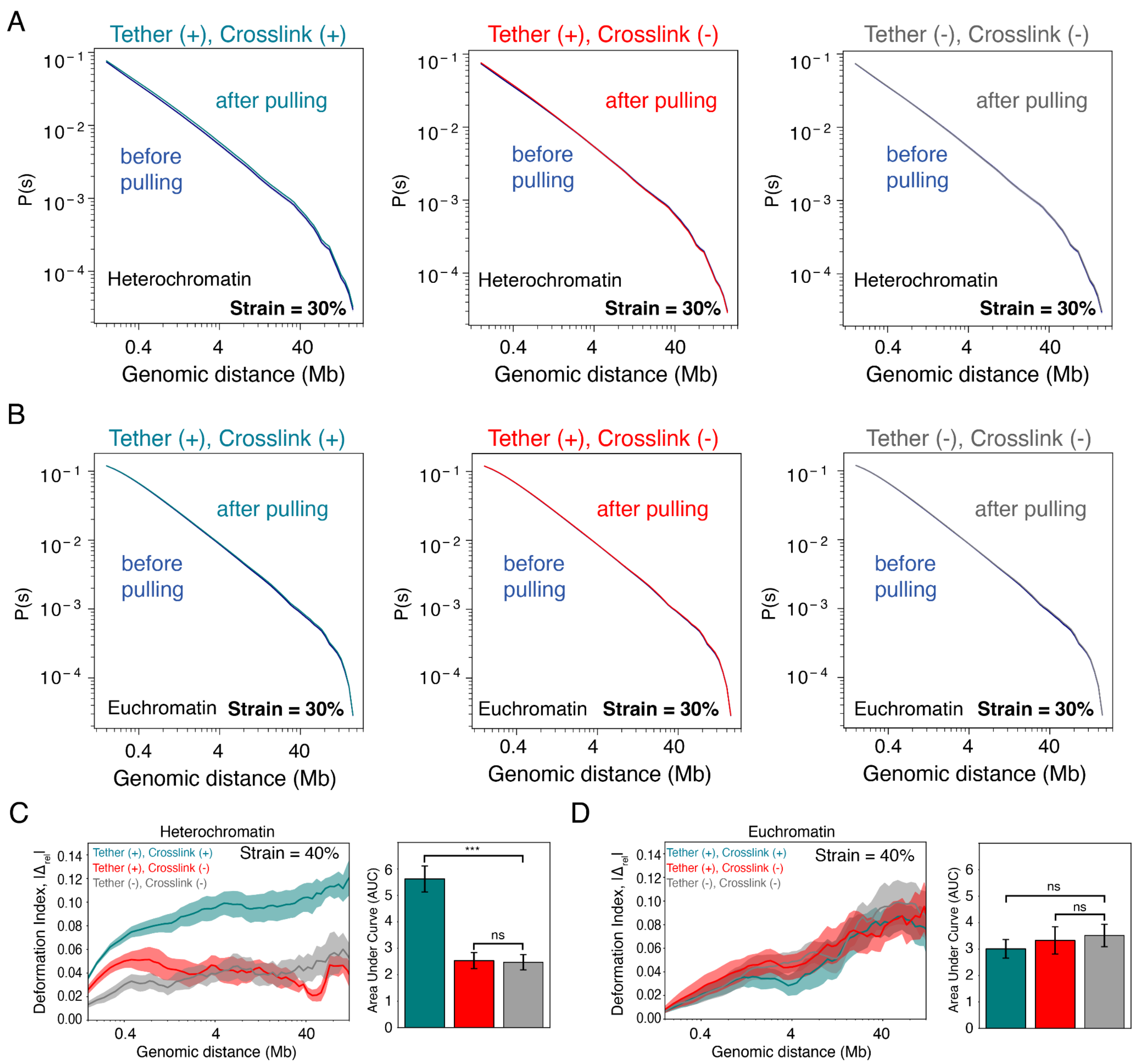


**Figure 14** Chromatin reorganization during nuclear deformation is regulated by peripheral facultative heterochromatin-lamina tethering and crosslinking. A) Chromatin contact frequencies as a function of genomic distance for intra-facultative heterochromatin contacts for the genomic distances of 4-40 Mb, before and after 30% nuclear strain for nuclei with both lamina tethering and heterochromatin crosslinking (left), tethering but no crosslinking (center), or neither tethering nor crosslinking (right). B) Chromatin contact frequencies as a function of genomic distance for intra-euchromatin contacts for the genomic distances of 4-40 Mb, before and after 30% nuclear strain for nuclei with both lamina tethering and heterochromatin crosslinking (left), tethering but no crosslinking (center), or neither tethering nor crosslinking (right). C) Absolute relative change, |Δrel|, in heterochromatic contacts as a function of genomic distance after deformation of nuclei with both lamina tethering and heterochromatin crosslinking (turquoise), tethering but no crosslinking (red), or neither tethering nor crosslinking (gray). Right: Bar plot showing the total absolute relative change, summed over all genomic distances. D) Absolute

relative change, |Δrel|, in euchromatic contacts as a function of genomic distance after deformation of nuclei with both lamina tethering and heterochromatin crosslinking (turquoise), tethering but no crosslinking (red), or neither tethering nor crosslinking (gray). Right: Bar plot showing the total absolute relative change, summed over all genomic distances. Bars are mean ± SEM. Statistical significance is assessed by one-way ANOVA followed by a post-hoc Tukey's HSD test for pairwise comparisons. * denotes $p < 0.05$, ** denotes $p < 0.01$, and *** denotes $p < 0.001$. n=10.

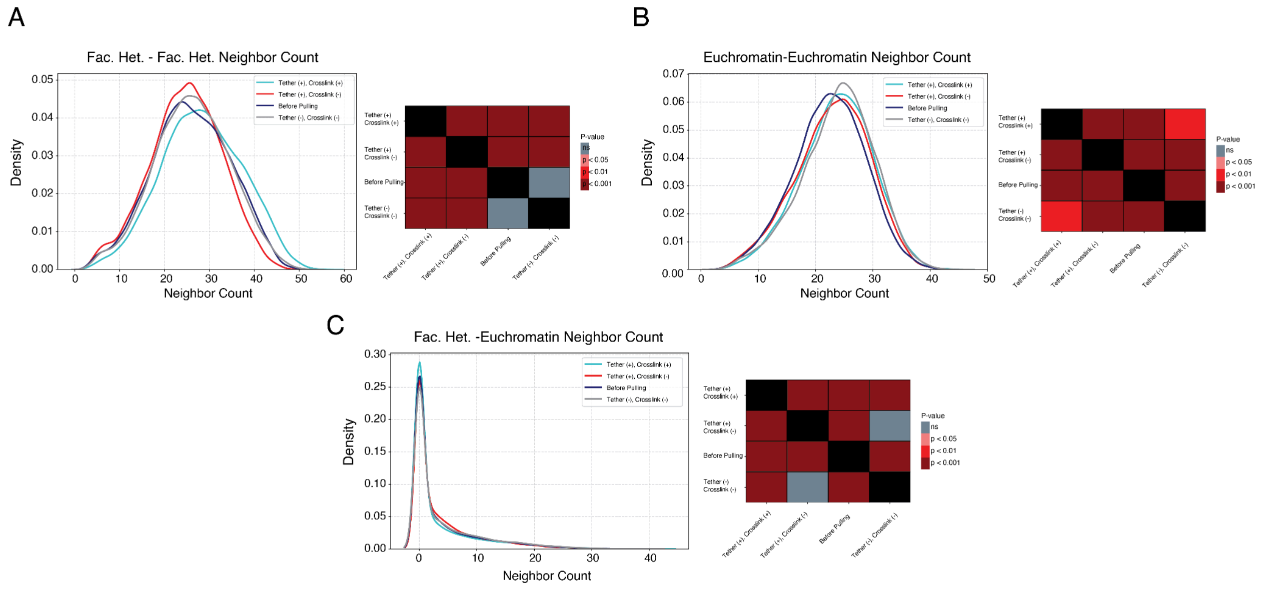


**Figure 15.** Chromatin reorganization during nuclear deformation is regulated by peripheral facultative heterochromatin-lamina tethering and crosslinking. A) Kernel density plot demonstrating the distribution of the facultative heterochromatin - facultative heterochromatin contacts including both intra and inter-chromatin contacts. B) Kernel density plot demonstrating the distribution of the euchromatin-euchromatin contacts including both intra and inter-chromatin contacts. Heatmap represents the p values from multiple comparisons through Kruskal-wallis followed by post hoc dunn. C) Kernel density plot demonstrating the distribution of the facultative heterochromatin-euchromatin contacts including both intra and inter-chromatin contacts. Heatmaps in, A, B and C, represent the p-values from multiple comparisons using the Kruskal-Wallis test followed by post hoc Dunn's test. n=10.
